# Inference of complex population histories using whole-genome sequences from multiple populations

**DOI:** 10.1101/026591

**Authors:** Matthias Steinrücken, Jack Kamm, Jeffrey P. Spence, Yun S. Song

## Abstract

There has been much interest in analyzing genome-scale DNA sequence data to infer population histories, but inference methods developed hitherto are limited in model complexity and computational scalability. Here we present an efficient, flexible statistical method, diCal2, that can utilize whole-genome sequence data from multiple populations to infer complex demographic models involving population size changes, population splits, admixture, and migration. Applying our method to data from Australian, East Asian, European, and Papuan populations, we find that the population ancestral to Australians and Papuans started separating from East Asians and Europeans about 100,000 years ago, and that the separation of East Asians and Europeans started about 50,000 years ago, with pervasive gene flow between all pairs of populations.

## INTRODUCTION

Whole-genome sequences are now routinely available for population genetic analyses, and inference methods that can take better advantage of genome-scale data have received considerable attention in recent years. In particular, there has been much interest in methods that can use the genomic data of individuals from multiple populations to infer complex models of population history. In addition to being of historical interest, population demography is important to study because it influences patterns of genetic variation, and understanding the intricate interplay between demography and other evolutionary forces such as natural selection is a major aim in population genetics.

Inferring these demographic histories is computationally and statistically challenging. One class of methods [1–8] based on the sample frequency spectrum (SFS) is computationally efficient but ignores linkage information, and the minimax rate of convergence for such estimators is poor [9]. Also, their utility is limited by the fact that the number of model parameters that is theoretically possible to estimate using the SFS alone is bounded by the sample size [10]. Methods [11–18] that take linkage structure into account are empirically more statistically efficient and can be used to infer models with many parameters even when the sample size is small. This is of practical importance, since an increasing number of studies now seek to infer complex demographic models involving multiple populations using only a small number of individuals sampled from each population (e.g., [19–22]). A popular demographic inference method of this kind is `PSMC` [11], which uses a pair of sequences to infer piecewise-constant population size histories. Its extension, `MSMC` [16], can use sequences sampled from a pair of populations to infer a genetic separation history in addition to population size changes. A more recent method called `smc++` [17] can scale to hundreds of individuals, but it is able to analyze individuals from only a pair of populations that have diverged without subsequent gene flow.

Parallel to these developments, an inference method called `diCal` (Demographic Inference using Composite Approximate Likelihood) [14] was introduced to infer piecewise-constant effective population size histories using multiple sequences, thereby providing improved inference about the recent past. The key mathematical component of `diCal` is the conditional sampling distribution (CSD) *π*_Θ_, which describes the conditional probability of observing a new sequence or haplotype given a collection of already observed haplotypes, under a given population genetic model with parameters Θ. The corresponding genealogical process can be formulated as a hidden Markov model (HMM), enabling efficient inference.

In this paper, we extend `diCal` in several ways to develop a scalable inference tool for population genomic analysis under general demographic models. To handle gene flow between populations, our new method, called `diCal2`, builds on previous theoretical work [23] which introduced a CSD for subdivided populations with unchanging continuous migration; that earlier work did not address parameter estimation, which is the focus of this article. In contrast to `MSMC`, which does not explicitly model population structure, we consider fully parametric demographic models, including subdivided population structure with migration that are easier to interpret. Our method also enables inference under demographic models more general than the two-population clean-split model currently implemented in `smc++`. Specifically, our method is flexible enough to model:

1. An arbitrary number of populations specified by the user.
2. An arbitrary pattern of population splits and mergers.
3. More general population size changes (e.g., piecewise-exponential).
4. Arbitrary migration patterns with time-varying continuous migration rates or pulse admixture events.
5. An arbitrary poly-allelic mutation model at each site (including di- or tetra-allelic).

These are significant improvements on the previous version of `diCal` [14], which could only be used to infer piecewise constant population size changes in a single population.

In addition to these features, we introduce major computational improvements which enable the use of whole-genome data. The mathematical details of our method and the computational extensions are provided in **Materials and Methods** and SI Appendix, SI Text. Below we briefly highlight the key technical advances.

In `PSMC` and the earlier version of `diCal`, the demographic epochs and HMM discretization intervals are both fixed, and the latter forms a strict refinement of the former. In contrast, discretization intervals and demographic epochs are decoupled in our improved version of `diCal`. For example, population size change-points or population split times can vary freely and do not need to coincide with discretization interval boundaries. This flexibility allows for more accurate parameter estimation, especially for population split times.

Moreover, the CSDs for different haplotypes can be combined in various ways to devise a composite likelihood that can be used in a maximum likelihood framework for parameter estimation. Our implementation of the expectation-maximization (EM) algorithm allows any composite likelihood that is composed of sums and products of CSDs, which includes the Product of Approximate Conditionals (PAC) used by Li & Stephens [24] to detect recombination hotspots.

For substantial computational speedup, we implement a previously described “locus-skipping” algorithm [25], which analytically and exactly integrates over contiguous stretches of non-segregating loci. However, locus-skipping is less computationally efficient with missing data, and thus, missing alleles should be imputed to take full advantage of the locus-skipping algorithm. To this end, we implement an alternative speedup by grouping loci together into larger blocks. A similar speedup was used in `PSMC`, but it treats the whole block as a single di-allelic site. In contrast, the blocks in our method are viewed as full non-recombining haplotypes.

For complex demographic models, the likelihood function may have local optima. To address this issue, we implement a flexible genetic algorithm and combine it with the EM procedure to enable more efficient navigation of high-dimensional parameter space.

We applied our method to data from the Simons Genome Diversity Project (SGDP) [20] to investigate the population history of Australians, East Asians, Europeans, and Papuans. There has been some debate whether the population ancestral to Australians and Papuans (which we call Australo-Papuans following [26]) split off prior to the divergence of East Asians and Europeans (e.g., [26]), or whether East Asians and Australo-Papuans first split from Europeans (e.g., [27]). We find substantial evidence in favor of the former hypothesis, but that there has been pervasive gene flow between all of these populations since their divergence.

## RESULTS

To demonstrate the flexibility, accuracy, and efficiency of our method, we performed an extensive simulation study under a variety of biologically relevant demographic scenarios. DNA sequence data were simulated using the software `scrm` [28]. We simulated 100 datasets for each demographic scenario and set the haplotype length to 250 Mbp for each dataset. We used 1.25 × 10^-8^ per generation for the per-site mutation and recombination rates and used a value of 100 kbp for the value of the `-l` parameter in `scrm`, which determines the length of the recombination history to be used during the simulation. Generation time was assumed to be 30 years. In each scenario, we used our method to estimate all demographic parameters of the underlying model.

### Recent exponential growth

The first model we investigated involves recent exponential population growth. To investigate the performance of our method, we simulated data consisting of 10 haplotypes under the demographic model depicted in Figure 1A. We fixed *T*_B_ = 65 ka, *T*_G_ = 15 ka, *N*_A_ = 15, 000, and *N*_B_ = 1,800, and used three different values for the growth rate *r*: 0.25%, 0.5%, and 1.0% per generation.

**Figure 1:**
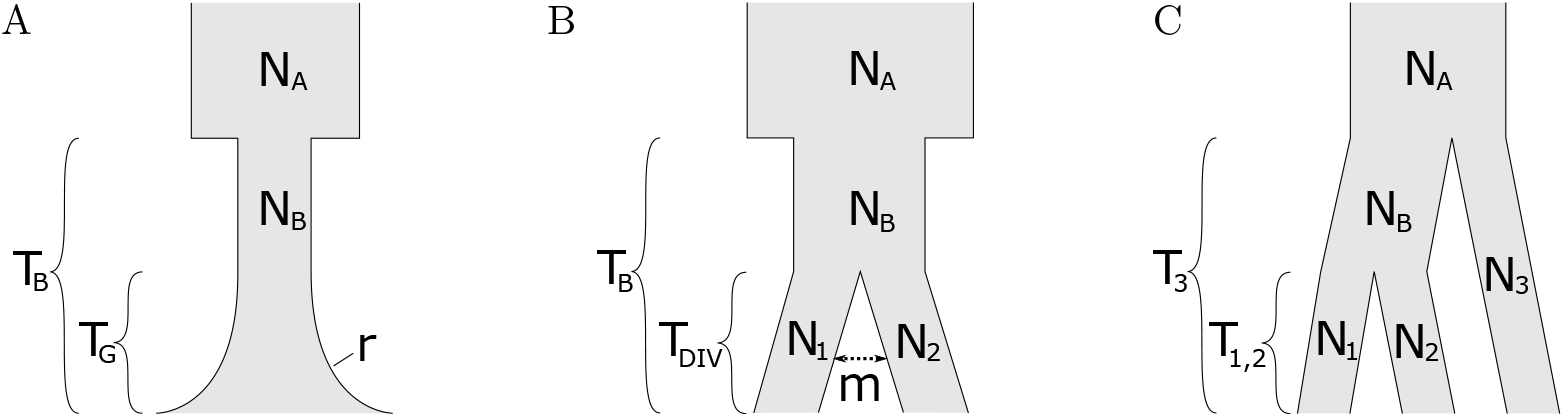
Demographic models used in our simulation study. (A) Recent exponential population growth: An ancestral population of size *N_A_* undergoes a bottleneck at time *T*_B_, where its size is reduced to *N*_B_. Growth starts at time *T*_G_ at an exponential rate *r*. (B) Demographic model of a population split: An ancestral population of size *N*_A_ undergoes a strong bottleneck that starts at time *T*_B_ in the past, and reduces the population size to *N*_B_. At time *T*_DIV_, this population then splits into two populations of size *N*_1_ and *N*_2_, respectively. Following the population split, migrants are exchanged at a rate *m*. (C) Demographic model of three populations with pure splits: An ancestral population of size *N_A_* splits into two populations of size *N_B_* and *N*_3_ at time *T*_3_. The former then again splits into two populations of size *N*_1_ and *N*_2_ at time *T*_1,2_.

We used the leave-one-out composite likelihood (LCL) in our EM procedure combined with a genetic algorithm to estimate all 5 parameters of the demographic model. For the genetic algorithm, we chose 50 random starting points that were each optimized for 5 EM iterations. Then we chose the 5 best parameter values (“parents”) and replaced each of them with an average of 3 “offspring” parameter sets to obtain the next “generation”. These were then optimized for 5 EM iterations. We repeated this procedure for 4 more “generations”, and reported the parameters that achieved the overall maximal likelihood value. We found that the results are robust to the choice of composite likelihood scheme.

Violin plots representing the accuracy of the inferences are shown in Figure 2A and Supplementary Figure S1. Analysis of the simulated data shows that in these scenarios, all parameters are estimated with little variability. However, the results indicate that the estimate of the exponential growth rate is biased upward. This bias is somewhat counterbalanced by a slight downward bias of the time when growth starts, and the population size before growth starts. In fact, the estimates lead to very accurate contemporary population sizes. We note that it is possible to empirically correct for biases in applications via simulation. Furthermore, using more sequence data for each individual reduces the variability of the estimates. We stress that our method accurately estimates recent exponential growth rates using only 10 haplotypes. This is far less than the sample size (thousands to tens of thousands) required by SFS-based methods to get reasonable estimates; see Bhaskar et al. [5] and references therein.

**Figure 2:**
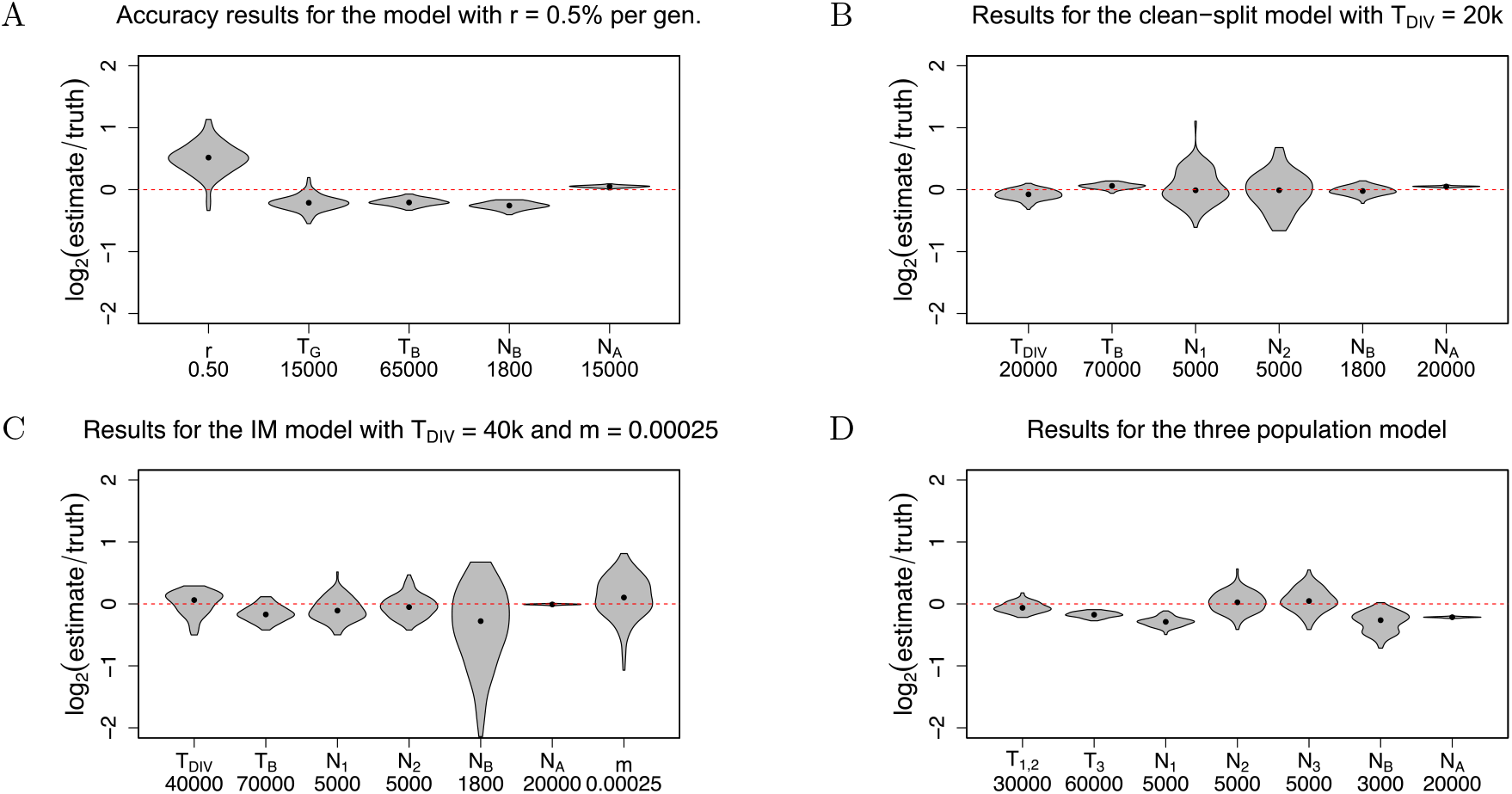
Accuracy results of our method, diCal2. Each violin plot shows the base-2 logarithm of the relative error (estimate/truth) for the analysis of 100 simulated datasets. Thus, a value of 0 corresponds to an exact estimate, whereas +1 is a two-fold over- and −1 is a two-fold underestimate. True parameter values are shown on the *x*-axis. (A) The recent exponential growth model shown in Figure 1A with expansion rate *r* = 0.5% per generation. Parameter estimates were obtained using only 10 haplotypes, which is much less than the sample size (thousands to tens of thousands) required by SFS-based methods to get good estimates. (B) Accuracy results for the clean-split model (no gene flow, *m* = 0) shown in Figure 1B with divergence time *T*_DIV_ = 20 ka. Using only two haplotypes in each extant population, the parameters of this clean-split model could be estimated very accurately. (C) Accuracy results for the isolation with migration (IM) model shown in Figure 1B with divergence time *T*_DIV_ = 40 ka, and migration probability *m* = 0.00025. As in the clean-split case, only two haplotypes in each extant population were used. Most parameter estimates show little bias or variability. See the text for further discussion. (D) Accuracy results for the three-population model shown in Figure 1C, using two haplotypes in each extant population.

### Population split

We also investigated a model of a past population split, depicted in Figure 1B. This model allows for a bottleneck before the populations split and subsequent gene flow following the split. We first focused on the case with no gene flow, i.e., with migration probability *m* = 0.

We simulated datasets with two haplotypes in each of the extant populations. We simulated 100 datasets each for *T*_DIV_ = 10 ka and 20 ka, with the remaining parameters set to *T*_B_ = 70,000 ya, *N*_A_ = 20, 000, *N*_B_ = 1, 800, and *N*_1_ = *N*_2_ = 5, 000. This scenario has recently been used in a study of the demographic history of Native Americans [19]. In addition, we simulated 100 datasets with *T*_DIV_ = 70 ka, setting *N*_B_ = *N*_A_ = 20,000, thereby also removing the need for *T*_B_. For the genetic algorithm, we used 60 random starting points, and 6 “parents” for each of the following 4 “generations” for the cases *T*_DIV_ = 10 ka and 20 ka, and 40 starting points and 5 “parents” for the case *T*_DIV_ = 70 ka. We used the LCL.

Figure 2B and Supplementary Figure S2 show the accuracy of the estimator. These empirical results demonstrate that our method is able to estimate the parameters in this clean-split model with high accuracy. Most parameters show little bias and the empirical distributions are very narrow. Only the estimates of the extant population sizes *N*_1_ and *N*_2_ for *T*_DIV_ = 10 ka and *T*_DIV_ = 20 ka show a somewhat higher variability. Since this time frame is very recent on an evolutionary timescale, either more sampled haplotypes or more sequence data are required to better estimate these parameters.

### Isolation with migration

We also investigated the demographic model shown in Figure 1B allowing for positive gene flow after the ancestral population splits into two. We set the migration probability to *m* = 0.00025; i.e., an individual from population 1 can have a parent from population 2, and vice versa, with a probability of 0.00025 per generation. Using this migration probability, we simulated 100 datasets each consisting of two haplotypes in each extant population, using *T*_DIV_ = 40 ka, *T*_B_ = 70 ka, *N*_A_ = 20, 000, *N*_B_ = 1,800, and *N*_1_ = *N*_2_ = 5,000. We also simulated 100 datasets using *T*_DIV_ = 70 ka, *N*_B_ = *N*_A_ = 20,000, and *N*_1_ = *N*_2_ = 5,000. In the former case we used 70 starting points and 6 “parents” for each “generation” in the genetic algorithm, whereas for the later we used 50 and 5, respectively.

Figure 2C and Supplementary Figure S3 show the accuracy of the estimator. In both scenarios, we used the pairwise composite likelihood (PCL). Again, most parameter estimates show little bias or variability, the exceptions being *N_B_* and *m* in the first scenario. However, we note that the evolutionary timescales involved are, again, rather short, and thus the number of events informative about these parameters is small. In practice, the variability could be reduced by using additional chromosomes.

### Three-population model

Lastly, we simulated data under the model depicted in Figure 1C relating three extant populations. Under this model, an ancestral population of size *N_A_* splits into two populations of size *N_B_* and *N*_3_ at time *T*_3_. The one of size *N_B_* then splits into two populations of size *N*_1_ and *N*_2_ at time *T*_1,2_. We simulated 100 datasets with two haplotypes in each of the extant populations. We set the parameters to *T*_1,2_ = 30 ka, *T*_3_ = 60 ka, *N_A_* = 20, 000, *N_B_* = 3, 000, and *N*_1_ = *N*_2_ = *N*_3_ = 5, 000. For the genetic algorithm, we chose 70 starting points, and 6 “parents”, and used the LCL. Figure 2D shows the accuracy of our method. Again, the empirical distribution of the estimates show little bias or variability. To our knowledge, our method is the first coalescent-HMM based method that can analyze a scenario with 3 extant populations each with more than one haplotype.

### Application to Simons Genome Diversity Project (SGDP) data

We used our method to investigate the pattern of population splits between Australians, East Asians, Europeans, and Papuans. There has been some debate about the relative ordering of population splits; specifically, there has been competing evidence about whether East Asians and Europeans split most recently (e.g., [26]) or whether Australo-Papuans and East Asians split most recently (e.g., [27]). To date these splits, we used Australian, French, Han, and Papuan individuals from the SGDP [20] and fit models for each of the six possible pairs of these populations, allowing for recent population size changes and pulse admixture. The model is depicted in Figure 3, and additional details are given in **Materials and Methods**. The estimates of the divergence time *T*_DIV_ and admixture fraction p together with confidence intervals obtained using a parametric bootstrapping approach are presented in Table 1. We found compelling evidence that Australo-Papuans and Eurasians diverged first, about 100 ka, with subsequent French-Han divergence at 53.6 ka, and the Australian-Papuan divergence at 33.9 ka. Note that while these estimates of divergence times are largely consistent with a tree, some estimates appear to imply slightly different split times (for example the Australian-Han divergence time is about 15 ka earlier than the Australian-French divergence time): this is likely due to model misspecification resulting from an overly simplistic model.

**Figure 3:**
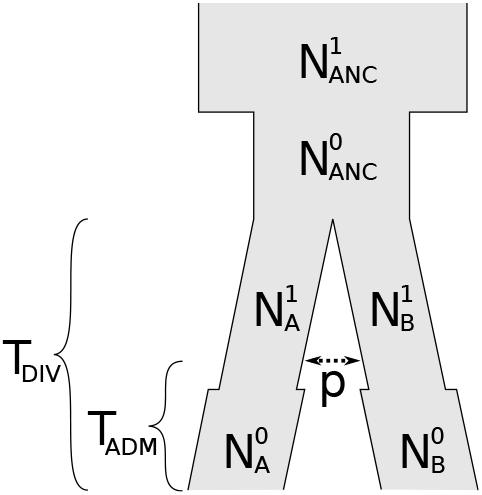
The demographic model used for the analysis of the French, Han, Papuan, and Australian population from the SGDP dataset. The ancestral population has two periods of constant size, then splits into two, and each of the extant populations has again 2 periods of constant size. Additionally, there is a symmetric pulse admixture event at *T*_ADM_, replacing *p*% of the ancestors in each population.

**Table 1:** Estimates of the divergence time *T*_DIV_ (in ka) and the admixture percentage *p* (in %) for the respective pair of populations from the SGDP dataset. The 95% confindence intervals obtained from the parametric bootstrap procedure are shown in square brackets.

|  | Han | Papuan | Australian |
| --- | --- | --- | --- |
| French | 54 ka [52,55]<br>14.8% [14.4,15.2] | 106 ka [104,108]<br>23.5% [23.0,24.0] | 106 ka [105,108]<br>25.0% [24.3,25.7] |
| Han |  | 113 ka [110,115]<br>26.3% [25.0,27.7] | 91 ka [91,91]<br>24.5% [23.7,25.4] |
| Papuan |  |  | 34 ka [33,35]<br>15.4% [14.6,16.1] |

We also found evidence of pervasive recent gene flow. In particular, we found pulse admixture proportions of 15-26% between each pair of populations all occurring 5-20 ka. We note that our model cannot capture all of the intricacies of human demographic history: there has likely been continuous gene flow between all populations punctuated by a few mass migrations. Our gene flow estimates likely attempt to capture both of these modes simultaneously along with indirect gene flow through intervening populations. While it is unlikely that about a quarter of any population was replaced by a geographically distant population, our results suggest that since their divergences tens of thousands of years ago, these populations have exchanged a considerable number of migrants.

Additional details about the data, data processing, and parameter settings for our method are presented in **Materials and Methods**. All parameter estimates, bootstrap results, and measures of goodness-of-fit evaluated using cross-coalescence rate curves may be found in Supplementary Table S1 and Supplementary Figure S4.

## DISCUSSION

The results described above demonstrate that our method can efficiently and accurately estimate demographic parameters in biologically relevant scenarios. Our method has recently been used to study the history of Native American peoples [19,21,22] due to the flexible framework underlying the method enabling the consideration of a wide range of population histories. Aside from demographic inference, we note that our method can be utilized in other population genetic problems of interest, such as model selection (see, for example, Supplementary Information 18.4 of [21]). Furthermore, the posterior decoding of latent variables in our CSD can be used in detecting admixture tracts [29], estimating fine-scale recombination rates in admixed individuals, distinguishing ancestral and introgressed polymorphism, and detecting incomplete lineage sorting. Also, applying our CSD in methods for phasing genotypes, imputing missing sequence data, and detecting identity-by-descent tracts [30] would make it possible to properly account for demography, thus potentially improving accuracy. Lastly, it is straightforward to incorporate temporal samples (ancient DNA sequences) into our method [21], which leads to further interesting applications.

## MATERIALS AND METHODS

Here, we briefly describe our method, a composite likelihood framework to estimate demographic parameters using EM. Further details are provided in SI Appendix, SI Text. We also describe our analysis of the SGDP data.

### Demographic inference using diCal2

A central building block of our novel method `diCal2` for demographic inference is the conditional sampling distribution (CSD) *π*_Θ_(*h*|*α*, **n**). It denotes the probability of observing the sequence or haplotype *h* in sub-population *α*, given that the haplotypes **n** have already been observed in their respective sub-populations and the underlying demography is described by the parameters Θ. The CSD can be described using a sequentially Markovian genealogical process [31,32] that approximates the true conditional genealogical process. Subsequent approximations to this sequential process lead to an HMM with finite hidden state space that can be used to efficiently compute approximate CSDs. We provide the details of the HMM approximations in Section 1 of Supporting Information Text. The CSDs presented in Steinrücken et. al. [23] and Sheehan et. al. [14] can be obtained as special cases of the model presented here.

The CSD can then be used to define composite likelihood functions, which in turn enables us to perform maximum composite likelihood inference of the demographic parameters Θ. We can use any such composite likelihood function that is composed of sums and products of CSDs, for example, the product of approximate conditionals (PAC) framework which has been used successfully by Li and Stephens [24] to infer recombination hotspots.

To find the parameter values that maximize this composite likelihood, we employ the composite likelihood in the standard expectation-maximization (EM) framework [33]. While in principle all parameters of the model can be inferred, we focus on the demographic parameters Θ. Since it is not possible to derive a closed form solution for the maximum in the maximization step in general, we employ numerical optimization schemes, like the Nelder-Mead simplex algorithm [34] to efficiently determine the requisite maximum. We provide mathematical details for the implementation of the EM algorithm in Section 3 of Supporting Information Text.

In Section 4 of Supporting Information Text, we provide details on the implementation of the “locus-skipping” algorithm, and the alternative speedup that groups loci into larger blocks. Furthermore, in Section 5 of Supporting Information Text, we provide mathematical details of the modifications to the trunk genealogy to increase accuracy. Finally, we describe in Section 6 of Supporting Information Text how to employ a discretization for the HMM computations that differs from the partition induced by the demography and remains fixed throughout the optimization procedure.

### Runtime

The runtime of the basic EM is linear in the number of haplotypes times the number of CSDs in the composite likelihood and quadratic in the number of populations involved. The E-step depends linearly on the length of the haplotypes, whereas the M-step is independent of this quantity. The exact complexity and runtime of parameter estimation depends on the composite likelihood used, the details of the genetic algorithm, and the number of parameters to estimate. The analyses of the simulated data presented in this section were performed on a cluster of AMD Opteron processors. The raw sequential runtime of analyzing a single dataset averaged 100–120 CPU-hours, but by taking advantage of the independence structure of the composite likelihood and the genetic algorithm, we were able to decrease the runtime to an average of 15–20 wall clock hours, using up to 16 cores in parallel. The one exception was the Three-population model, where the parallelized version took on average 70 wall clock hours, due to the more complex demographic model and the increased number of haplotypes.

### SGDP analysis

For the analysis of the SGDP data we used the following individuals: B_Australian-3, B_Australian-4, S_French-1, S_French-2, B_French-3, S_Han-1, S_Han-2, B_Han-3, S_Papuan-1, S_Papuan-3, and B_Papuan-15. The data were phased using `Shapeit` [35] with read-based phasing (phased data provided by I. Mathieson), and all sites in Heng Li’s 75bp universal mask [20] were treated as missing. As in our simulations, we used a mutation rate and recombination rate of 1.25 × 10^-8^ per-base per generation, and assumed a generation time of 30 years. For each pair of populations, we used all of the individuals from those populations and used all of the autosomes to fit a model where each population has a constant size from present to 5 ka, and another constant size from 5 ka until the divergence time of the populations. The ancestral population is assumed to be a constant size from the divergence time until 100 ka, beyond which we infer a separate constant size. We allow a symmetric pulse migration between the two populations. To fit this model, we performed 4 iterations of the genetic algorithm, starting from 15 arbitrary points, keeping the 3 best particles at each iteration, and then spawning a total of 10 particles. Each genetic algorithm iteration consisted of 6 EM iterations for each particle, using the LCL. For computational efficiency, we grouped loci into 2.5 kbp bins, and discretized time with 8 log-uniformly spaced break points between 1.5 ka and 5 Ma.

As seen in the simulations, the raw inferred parameters may be biased. To address this issue and infer confidence intervals, we performed a parametric bootstrap using msprime [36]. For each pair of populations, we simulated 10 full genome datasets, and re-ran our method on each of these datasets. Our reported estimates are “de-biased” estimates, obtained by subtracting the estimated bias from our raw estimates. We then used the bootstraps to estimate a standard deviation for each parameter, and reported confidence intervals based on a normal approximation (i.e. the de-biased estimate ±1.96 standard deviations) To avoid having these de-biased estimates or confidence intervals fall outside of the domain of the parameters (e.g. negative population sizes, times, or pulse proportions or pulse proportions > 100%), all de-biasing and confidence intervals were computed in log-space for population sizes and times and in logit-space for pulse proportions. The resulting estimates and confidence intervals were then transformed back to their natural space using the exponential map and logistic map, respectively. We note that this procedure means that our estimates are unbiased in log-space or logit-space, and may be slightly biased in their natural scale. All parameter estimates and bootstrap results are presented in Supplementary Table S1.

To assess goodness of fit, we used MSMC to infer cross-coalescence rate curves (CCRs) on the real data, and then on data simulated under our de-biased estimates, presented in Supplementary Figure S4. For each pair of populations, we used a single diploid from each population (B_Australian-3, S_French-1, S_Han-1, and S_Papuan-1), using all of the autosomes and again treating all sites in Heng Li’s 75bp universal mask as missing. We simulated 5 replicates of each pair of populations to assess the variability in the `MSMC` CCRs. The CCRs are qualitatively similar between the real and simulated data, and the fit is quite good for a model with only 9 parameters. As discussed above, introducing additional size changes and migration rates would likely improve the fit.

### Software availability

The algorithms described here are implemented in a new version of the software package `diCal2`, which is available for download at https://sourceforge.net/projects/dical2.

## Acknowledgments

We thank Sara Mathieson and Geno Guerra for helpful discussions and for testing our software. Furthermore, we thank Iain Mathieson for helpful discussions and providing the phased SGDP data. This research is supported in part by an NIH Grant R01-GM094402, and a Packard Fellowship for Science and Engineering. YSS is a Chan Zuckerberg Biohub Investigator.

