## Supplementary Information for "Inference of complex population histories using whole-genome sequences from multiple populations"

#### This PDF file includes:

Supporting Information Text
Figures S1 to S6
Table S1
References for Supplementary Information

### Supporting Information Text

### 1. Hidden Markov Model formulation of the approximate CSDs

In this section, we present some additional notation and results to describe the Hidden Markov Model (HMM) that can be used to approximate the conditional sampling distribution (CSD).

**1.1. Notation.** We model the sampled haplotypes under the finite-sites, finite-alleles coalescent with recombination. Denote the set of possible alleles at a specific site or locus by $E$. A haplotype $h$ of length $L$ carries an allele at every locus and is thus an $L$-tuple from the set $\mathcal{H} = E^L$ of possible haplotypes. Denote by $h[l]$ the allele that haplotype $h$ carries at locus $l$, and by $h[l : l']$ the vector $(h[l], \ldots, h[l'])$. At each locus, mutations can occur at a coalescent-scaled per-locus mutation rate of $\theta/2$, where $\theta = 4N_0\mu$, with $N_0$ being the reference effective population size and $\mu$ the per-locus per-generation mutation probability. Denote by $P$ the stochastic mutation matrix, that is, if a mutation occurs, then allele $a$ mutates into allele $a'$ with probability $P_{a,a'}$, for $a, a' \in E$. A crossover recombination event occurs between each pair of consecutive loci $(l, l+1)$, for $1 \leq l < L$, at coalescent-scaled rate of $\rho/2$, where $\rho = 4N_0 r$ and $r$ denotes the per-generation recombination probability. The recombination and mutation rates could, in principle, vary along the sequence, but for notational convenience we will assume that the rates are constant.

We assume that the haplotypes are sampled in any of $g$ extant populations, and denote the set of possible populations at present by $\Gamma = \{1, \ldots, g\}$. A sample configuration $\mathbf{n}$ can be described by a collection of non-negative integers $n_{\gamma,h} \geq 0$, which give the number of haplotypes of type $h \in \mathcal{H}$ sampled in population $\gamma \in \Gamma$. The total number of sampled haplotypes is denoted by $n = \sum_{\gamma\in\Gamma} \sum_{h\in\mathcal{H}} n_{\gamma,h}$. Further, $\mathbf{n}_\gamma$ denotes the configuration consisting of only those haplotypes sampled in population $\gamma$, and $n_\gamma = \sum_{h\in\mathcal{H}} n_{\gamma,h}$ denotes the number of such haplotypes.

We allow for a general demographic model, where the demographic structure and the migration rates can differ at different times in the past. To this end, choose $E + 1$ times $0 = t_0 \leq t_1 \leq \cdots \leq t_E = \infty$ to obtain a partition of the positive real line $[0, \infty)$ into $E$ epochs denoted by $I_\epsilon = [t_{\epsilon-1}, t_\epsilon)$. Here $t_0 = 0$ corresponds to the present and $t_E = \infty$ to an infinite time in the past. Denote the set of all epochs by $\mathcal{E} := \{1, \ldots, E\}$. Note that this notation allows for an epoch to have length zero. To allow for changes in the ancient demographic structure, define for each epoch $\epsilon \in \mathcal{E}$ a partition $\Gamma_\epsilon = \{\gamma_\epsilon^{(1)}, \ldots, \gamma_\epsilon^{(g_\epsilon)}\}$ of $\Gamma$, where all present populations whose indices are in the set $\gamma_\epsilon^{(i)}$ derive from the $i$th ancestral population in epoch $\epsilon$. Thus, there are $g_\epsilon = |\Gamma_\epsilon|$ populations during that epoch. We require that $\gamma_1^{(i)} = \{i\}, \forall i$ in the first epoch, $\Gamma_1 = \{\gamma_1^{(1)}, \ldots, \gamma_1^{(g)}\}$, and that for all $\epsilon \in \mathcal{E}\backslash\{E\}$ the partition $\Gamma_\epsilon$ is a refinement of $\Gamma_{\epsilon+1}$.

The size of population $\gamma \in \Gamma_\epsilon$ is given by $\kappa_\gamma^{(\epsilon)} N_0$, and the coalescent rate is inversely proportional to the population size. Furthermore, during an epoch $\epsilon$ of positive length, migration (backwards in time) from population $\gamma \in \Gamma_\epsilon$ into population $\delta \in \Gamma_\epsilon$ occurs at a coalescent-scaled rate of $m_{\gamma,\delta}^{(\epsilon)}/2$. Here $m_{\gamma,\delta}^{(\epsilon)} = 4N_0 v_{\gamma,\delta}^{(\epsilon)}$ and $v_{\gamma,\delta}^{(\epsilon)}$ is the per-generation probability that an individual in population $\gamma$ has a parent from population $\delta$, also known as the backward migration rate. To handle scenarios of population admixture we introduce a mechanism for instantaneous migration during an epoch $\epsilon$ of length zero, where $t_{\epsilon-1} = t_\epsilon$ and $I_\epsilon = \emptyset$ hold. Instantaneous migration from population $\gamma$ to $\delta$ during such an epoch occurs with probability $y_{\gamma,\delta}^{(\epsilon)}$, the probability that an individual residing in population $\gamma \in \Gamma_\epsilon$ at time $t_{\epsilon-1}$ has an ancestor residing in population $\delta \in \Gamma_\epsilon$ at time $t_\epsilon$. We denote all the parameters necessary to describe a demographic history by $\Theta$, and present an example in Figure S5.

**1.2. Demography-aware CSD using Trunk Approximation.** Recall that the CSD $\pi_\Theta(h|\alpha, \mathbf{n})$ denotes the probability of observing the haplotype $h$ in sub-population $\alpha$, given that the haplotypes $\mathbf{n}$ have already been observed and the underlying demography is described by the parameters $\Theta$. Computing the true CSD $\pi_\Theta(h|\alpha, \mathbf{n})$ requires integrating over all possible genealogies relating the haplotypes in the already observed configuration $\mathbf{n}$ and the possible ways of attaching the lineage of the additional haplotype $h$ to these genealogies. To approximate this high-dimensional integral, assume that the unknown genealogy of the configuration $\mathbf{n}$ is given by an unchanging "trunk" of ancestral lineages for each haplotype extending infinitely into the past. If populations are merged at some point in the past, then the trunk-lineage continues in the merged population. Paul and Song (1) and Steinrücken et al. (2) motivated this approximation using an approach based on the generator of the underlying diffusion process (3, 4), and provided an extensive analysis of its accuracy. The trunk-approximation for a given configuration $\mathbf{n}$ is depicted in Figure S6A.

The following generative process describes the distribution of the ancestral lineage and the allelic composition of the additional haplotype $H$ under the trunk approximation $\pi^T_\Theta(\cdot|\alpha, \mathbf{n})$. First, a sequence of marginal additional ancestral lineages is sampled that include, at each locus, a history of migration events performed by ancestors of $H$ along the ancestral lineage at this locus, the lineage of the trunk into which the ancestor coalesces, and the times of these events. Under assumptions similar to the Sequentially Markov Coalescent (5, 6), these marginal lineages can be generated sequentially starting from the first (left-most) locus in a Markovian fashion. At the first locus, an additional ancestral lineage starts at the present in population $\alpha$ and extends into the past. During an epoch $\epsilon$ of positive length, if the lineage resides in population $\gamma \in \Gamma_\epsilon$, then it is subject to the events:

- *Migration:* The lineage migrates to population $\delta \in \Gamma_\epsilon$ with rate $m^{(\epsilon)}_{\gamma,\delta}$.
- *Absorption:* The lineage is absorbed into a uniformly chosen trunk-lineage in the population it currently resides in at rate $\left(\kappa^{(\epsilon)}_\gamma\right)^{-1}$, the inverse of its size.

During an epoch $\epsilon$ of length zero, the only possible event is

- *Pulse-migration:* The additional lineage migrates to population $\delta \in \Gamma_\epsilon$ with probability $y^{(\epsilon)}_{\gamma,\delta}$.

If at the end of an epoch $\epsilon$ the additional lineage resides in a population that merges with other populations into a single ancestral population in epoch $\epsilon + 1$, then it continues in the ancestral population after time $t_\epsilon$. This Markov process specifies the initial distribution at the first locus and also describes the marginal distribution of an additional lineage. The migration rate is two-fold higher then in the standard coalescent to balance out the non-migrating trunk.

Under the full coalescent with recombination, the ancestral lineages of two loci that are separated by a recombination distance $\rho$ evolve together into the past for an exponentially distributed amount of time with parameter $\rho/2$ until they are decoupled by a recombination event, and evolve independently beyond this event. Thus, under the approximate CSD $\pi^T_\Theta(\cdot|\alpha, \mathbf{n})$, given the marginal additional genealogy at a certain locus $l-1$, the marginal additional genealogy at locus $l$ is sampled as follows. Denote the time of absorption at locus $l-1$ by $t_{l-1}$. To determine whether and at what time an ancestral recombination event separates locus $l-1$ and locus $l$, a time $t_b$ is sampled from an exponential distribution with parameter $\rho$. If $t_b > t_{l-1}$, then the two loci are not separated by an ancestral recombination event. In this case, the complete marginal additional genealogy at locus $l-1$ is copied to the next locus, including the history of migration events, thus $t_{l-1} = t_l$. If $t_b \leq t_{l-1}$, then a recombination event separates the two loci. In this case, the marginal additional lineage at locus $l-1$ from the present up to the time of the breakpoint $t_b$ is copied to locus $l$, including the population it resides in at that time. The marginal additional lineage at locus $l$ beyond the time of the breakpoint then evolves independently according to the marginal dynamics, that is, it is independently subject to the migration dynamics until it is ultimately absorbed into a lineage of the trunk. Note that the recombination rate is two-fold higher than in the standard coalescent to again compensate for the lack of events in the trunk.

Once a sequence of marginal additional genealogies is generated, the alleles of the additional haplotype are sampled as follows. At each locus, the allele carried by the haplotype corresponding to the absorbing lineage at the respective locus is propagated along the marginal additional lineage of length $t_l$ from the time of absorption to the present. Mutation events occur at rate $\theta$ and change the current allele according to the stochastic mutation matrix $P$. Note that the rate of evolution is again multiplied by two. This generative process describes the distribution of the additional haplotype under $\pi^T_\Theta(\cdot|\alpha, \mathbf{n})$. A realization can be seen in Figure S6A.

**1.3. Markov chain governing the marginal dynamics.** We now introduce the mathematical notation to formalize the backward in time Markov chain that governs the marginal migration and absorption dynamics in our CSD. Furthermore, we provide details on how to compute the requisite transition probabilities for this Markov chain.

***1.3.1. Migration matrices.*** The migration rates for a given epoch $\epsilon$ can be subsumed in the migration matrix

$$M_\epsilon := \begin{pmatrix} -m_1^{(\epsilon)} & m_{1,2}^{(\epsilon)} & \cdots & m_{1,g_\epsilon}^{(\epsilon)} \\ m_{2,1}^{(\epsilon)} & -m_2^{(\epsilon)} & \cdots & m_{2,g_\epsilon}^{(\epsilon)} \\ \vdots & \vdots & \ddots & \vdots \\ m_{g_\epsilon,1}^{(\epsilon)} & \cdots & \cdots & -m_{g_\epsilon}^{(\epsilon)} \end{pmatrix}, \quad [1]$$

where we denoted the elements in $\Gamma_\epsilon$ by $1, \ldots, g_\epsilon$, and $m_\gamma^{(\epsilon)} = \sum_{\delta \neq \gamma} m_{\gamma,\delta}^{(\epsilon)}$ for each $\gamma \in \Gamma_\epsilon$.

Along similar lines, for epochs of length zero with $t_{\epsilon-1} = t_\epsilon$, the matrix

$$Y_\epsilon := \begin{pmatrix} y_{1,1}^{(\epsilon)} & y_{1,2}^{(\epsilon)} & \cdots & y_{1,g_\epsilon}^{(\epsilon)} \\ y_{2,1}^{(\epsilon)} & y_{2,2}^{(\epsilon)} & \cdots & y_{2,g_\epsilon}^{(\epsilon)} \\ \vdots & \vdots & \ddots & \vdots \\ y_{g_\epsilon,1}^{(\epsilon)} & \cdots & \cdots & y_{g_\epsilon,g_\epsilon}^{(\epsilon)} \end{pmatrix} \quad [2]$$

comprises the instantaneous migration probabilities.

***1.3.2. Extended migration matrix.*** As described in Section 1.2, during an epoch $\epsilon$ of positive length, in addition to the migration dynamics described by the migration matrix defined in (1), the marginal additional lineage can be absorbed into a lineage of the trunk in the sub-population it currently resides in.

To model this behavior, the Markov chain describing the dynamics has two states per sub-population. One state for the case when the lineage only resides in the respective sub-population, and one state for the case when the lineage is actually absorbed. The dynamics between the unabsorbed states is governed by the migration rates given in the migration matrix $M_\epsilon$. An absorbed state can only be reached from the unabsorbed state associated with the same sub-population, since the additional lineage can only be absorbed into a trunk-lineage in the sub-population it currently resides in. While the lineage resides in sub-population $\gamma \in \Gamma_\epsilon$, it gets absorbed with rate $(\kappa_\gamma^{(\epsilon)})^{-1} n_\gamma$, proportional to the inverse population size $(\kappa_\gamma^{(\epsilon)})^{-1}$ and $n_\gamma$, the number of trunk lineages in the sub-population. The latter is given as $n_\gamma = \sum_{\delta \in \gamma} n_\delta$, the sum over the number of haplotypes in all present sub-populations that $\gamma$ is ancestral to. Furthermore, since the Markov chain cannot exit an absorbed state, the rates for leaving absorbed states are zero.

Thus, the Markov chain describing the migration and absorption dynamics for epoch $\epsilon$ backwards in time evolves according to the $2g_\epsilon \times 2g_\epsilon$ rate matrix

$$Z_\epsilon := \begin{pmatrix} M_\epsilon - A_\epsilon & A_\epsilon \\ 0 & 0, \end{pmatrix} \quad [3]$$

where the matrix

$$A_\epsilon = \mathrm{diag}\Big((\kappa_{\gamma_\epsilon^{(1)}}^{(\epsilon)})^{-1} n_{\gamma_1}, \ldots, (\kappa_{\gamma_\epsilon^{(g)}}^{(\epsilon)})^{-1} n_{\gamma_g}\Big), \quad [4]$$

for $\gamma_\epsilon^{(i)} \in \Gamma_\epsilon$ and $g = |\Gamma_\epsilon|$, governs the absorption of the additional lineage into the trunk. Further, let $a_{\gamma_\epsilon^{(i)}}$ denote the index in this matrix of the state "being absorbed in $\gamma_\epsilon^{(i)}$."

*1.3.3. Spectral representation (Eigendecomposition).* For an epoch $\epsilon$ of positive length, the spectral representation of $Z_\epsilon$ is helpful to compute certain integrals and matrix exponentials necessary for calculating the requisite probabilities of the HMM underlying the CSD $\pi^D_\Theta$. Assume that the $2g_\epsilon \times 2g_\epsilon$ matrix $Z_\epsilon$ is diagonalizable, and denote by $\{\lambda^{(\epsilon)}_1, \dots \lambda^{(\epsilon)}_{2g_\epsilon}\}$ the eigenvalues and by $\{v^{(\epsilon)}_1, \dots, v^{(\epsilon)}_{2g_\epsilon}\}$ the corresponding eigenvectors. Note that $g_\epsilon$ eigenvalues are zero due to the $g_\epsilon$ absorbing states.

Now define

$$V_\epsilon := \begin{pmatrix} v^{(\epsilon)}_1 & \cdots & v^{(\epsilon)}_{2g_\epsilon} \end{pmatrix} \qquad [5]$$

to be the matrix that has the eigenvectors as columns. With this definition we can write

$$Z_\epsilon = V_\epsilon \begin{pmatrix} \lambda^{(\epsilon)}_1 & \cdots & 0 \\ \vdots & \ddots & \vdots \\ 0 & \cdots & \lambda^{(\epsilon)}_{2g_\epsilon} \end{pmatrix} V^{-1}_\epsilon = \sum_{k=1}^{2g_\epsilon} \lambda^{(\epsilon)}_k v^{(\epsilon)}_k w^{(\epsilon)}_k, \qquad [6]$$

where $w^{(\epsilon)}_k$ is the $k$-th row of $V^{-1}_\epsilon$, which in turn yields

$$e^{tZ_\epsilon} = V_\epsilon \begin{pmatrix} e^{t\lambda^{(\epsilon)}_1} & \cdots & 0 \\ \vdots & \ddots & \vdots \\ 0 & \cdots & e^{t\lambda^{(\epsilon)}_{2g_\epsilon}} \end{pmatrix} V^{-1}_\epsilon = \sum_{k=1}^{2g_\epsilon} e^{t\lambda^{(\epsilon)}_k} v^{(\epsilon)}_k w^{(\epsilon)}_k. \qquad [7]$$

Then

$$\left(e^{tZ_\epsilon}\right)_{\gamma,\delta} = \sum_{k=1}^{2g_\epsilon} e^{t\lambda^{(\epsilon)}_k} (v^{(\epsilon)}_k w^{(\epsilon)}_k)_{\gamma,\delta} \qquad [8]$$

holds, and furthermore,

$$\left(Z_\epsilon e^{tZ_\epsilon}\right)_{\gamma,\delta} = \sum_{k=1}^{2g_\epsilon} \lambda^{(\epsilon)}_k e^{t\lambda^{(\epsilon)}_k} (v^{(\epsilon)}_k w^{(\epsilon)}_k)_{\gamma,\delta}. \qquad [9]$$

From equations (8) and (9) it follows that

$$\frac{d}{dt}\left(e^{tZ_\epsilon}\right)_{\gamma,\delta} = \left(Z_\epsilon e^{tZ_\epsilon}\right)_{\gamma,\delta}. \qquad [10]$$

Note that if $Z_\epsilon$ is not diagonalizable, a similar spectral decomposition could be employed, using generalized eigenvalues and the Jordan normal form. However, for ease of notation, we will only present the computations in the sequel for diagonalizable matrices.

**1.4. Continuous HMM.** We now introduce the initial, the transition, and the emission probability for the HMM with continuous absorption time, to illustrate our approach and introduce some useful concepts. At locus $l$, denote by $T^A_l$ the random absorption time, by $G_l$ the random population where the absorption event takes place, and by $X_l$ the random trunk-lineage that the additional lineage is absorbed into. Since lineages in the trunk do not migrate, the absorbing lineage $X_l$ would be sufficient to determine the population where absorption takes place. However, we keep the population explicit for later convenience.

*1.4.1. Marginal/Initial density.* The transition density in this model is reversible with respect to the initial density, thus the initial and marginal densities are identical. They can be obtained as follows.

First, define

$$f^\epsilon_{\mu_\epsilon,\gamma_\epsilon} := \begin{cases} \left(e^{(t_\epsilon - t_{\epsilon-1})Z_\epsilon}\right)_{\mu_\epsilon,\zeta_\epsilon}, & \text{if } I_\epsilon \neq \emptyset, \\ \left(Y_\epsilon\right)_{\mu_\epsilon,\zeta_\epsilon}, & \text{if } I_\epsilon = \emptyset, \end{cases} \qquad [11]$$

the probability that a lineage residing in sub-population $\mu_\epsilon \in \Gamma_\epsilon$ at time $t_{\epsilon-1}$ resides in sub-population $\gamma_\epsilon \in \Gamma_\epsilon$ at time $t_\epsilon$. In an epoch of length zero ($I_\epsilon = \emptyset$), this given by the instantaneous migration probabilities, whereas in an epoch of positive length ($I_\epsilon \neq \emptyset$), the matrix exponential of the extended migration matrix accounts for the fact that the lineage is not absorbed during the interval $I_\epsilon$.

The quantity (11) can be employed to recursively define the probability $p^{(0,\epsilon-1)}_{\alpha,\gamma_\epsilon}$ that the additional lineage resides in sub-population $\alpha \in \Gamma_1$ (where the additional haplotype is sampled) at the beginning of epoch 1 (time $t_0$) and resides in sub-population $\gamma_\epsilon \in \Gamma_\epsilon$ at $t_{\epsilon-1}$, while not having been absorbed by that time. The latter can thus be calculated by dynamic programming using the formulas $p^{0,0}_{\alpha,\gamma_1} = \delta_{\alpha,\gamma_1}$, where $\delta$ is the Kronecker-delta, and

$$p^{(0,\epsilon-1)}_{\alpha,\gamma_\epsilon} = \sum_{\mu_{\epsilon-1}\in\Gamma_{\epsilon-1}} \sum_{\substack{\zeta_{\epsilon-1}\in\Gamma_{\epsilon-1}\\ \zeta_{\epsilon-1}\subset\gamma_\epsilon}} p^{(0,\epsilon-2)}_{\alpha,\mu_{\epsilon-1}} f^{\epsilon-1}_{\mu_{\epsilon-1},\zeta_{\epsilon-1}}. \tag{12}$$

The sum $\sum_{\substack{\zeta_{\epsilon-1}\in\Gamma_{\epsilon-1}\\ \zeta_{\epsilon-1}\subset\gamma_\epsilon}}$ is necessary, since it sums over all the sub-populations that merge into the sub-population $\gamma_\epsilon$ at time $t_{\epsilon-1}$, and thus their probabilities have to be combined.

Now, for an arbitrary locus $l$ and a time $t_l \in \mathbb{R}_{\geq 0}$, let $e = \epsilon(t_l)$ denote the epoch of positive length such that $t_l \in I_e$. With $\omega_l \in \Gamma_e$, and $x_l \in \mathbf{n}_{\omega_l}$, the marginal density is then given as

$$\begin{aligned}\mathbb{P}\{T^A_l \in dt_l, G_l = \omega_l, X_l = x_l\} &= \frac{1}{n_{\omega_l}} \sum_{\gamma_e\in\Gamma_e} p^{(0,e-1)}_{\alpha,\gamma_e} \big(Z_e e^{(t_l - t_{e-1})Z_e}\big)_{\gamma_e,a_{\omega_l}} \\ &=: \frac{1}{n_{\omega_l}} q^{(0,e)}_{\alpha,a_{\omega_l}}(t_l - t_{e-1}).\end{aligned} \tag{13}$$

Here $\big(Z_e e^{(t_l - t_{e-1})Z_e}\big)_{\gamma_e,a_{\omega_l}}$ is the density of the event that the additional lineage is absorbed into a trunk-lineage in sub-population $\omega_l$ at time $t_l$. The factor $\frac{1}{n_{\omega_l}}$ appears, since it is absorbed into a specific trunk-lineage in this sub-population.

***1.4.2. Transition density.*** For ease of exposition, we focus on deriving the joint density first, which can then be combined with the marginal density to obtain the transition density. The additional lineages at two loci $l-1$ and $l$ can either be separated by a recombination event or not. The time of the recombination $T^B$ event is given by an exponential random variable with rate $\rho$.

Thus, with $t_{l-1}, t_l \in \mathbb{R}_{\geq 0}$, $\omega_{l-1} \in \Gamma_{\epsilon(t_{l-1})}$, $x_l \in \mathbf{n}_{\omega_{l-1}}$, $\omega_l \in \Gamma_{\epsilon(t_l)}$, and $x_l \in \mathbf{n}_{\omega_l}$, partitioning with respect to the time of the recombination event yields

$$\begin{aligned}&\mathbb{P}\{T^A_{l-1} \in dt_{l-1}, G_{l-1} = \omega_{l-1}, X_{l-1} = x_{l-1}, T^A_l \in dt_l, G_l = \omega_l, X_l = x_l\} \\ &\quad = \int_{t_b=0}^{\infty} \mathbb{P}\{T^A_{l-1} \in dt_{l-1}, G_{l-1} = \omega_{l-1}, X_{l-1} = x_{l-1}, T^A_l \in dt_l, G_l = \omega_l, X_l = x_l, T^B \in dt_b\} \\ &\quad = \int_{t_b=t_{l-1}\wedge t_l}^{\infty} \mathbb{P}\{T^A_{l-1} \in dt_{l-1}, G_{l-1} = \omega_{l-1}, X_{l-1} = x_{l-1}, T^A_l \in dt_l, G_l = \omega_l, X_l = x_l, T^B \in dt_b\} \\ &\qquad + \int_{t_b=0}^{t_{l-1}\wedge t_l} \mathbb{P}\{T^A_{l-1} \in dt_{l-1}, G_{l-1} = \omega_{l-1}, X_{l-1} = x_{l-1}, T^A_l \in dt_l, G_l = \omega_l, X_l = x_l, T^B \in dt_b\}\end{aligned} \tag{14}$$

for the joint distribution of the hidden states at locus $l-1$ and $l$.

The first term in (14) represents the case when the lineages at both loci are absorbed together before the recombination event can decouple them. It is given by

$$\begin{aligned}&\int_{t_b=t_{l-1}\wedge t_l}^{\infty} \mathbb{P}\{T^A_{l-1} \in dt_{l-1}, G_{l-1} = \omega_{l-1}, X_{l-1} = x_{l-1}, T^A_l \in dt_l, G_l = \omega_l, X_l = x_l, T^B \in dt_b\} \\ &\qquad = \delta_{t_{l-1},t_l}\delta_{\omega_{l-1},\omega_l}\delta_{x_{l-1},x_l} \frac{1}{n_{\omega_{l-1}}} q^{(0,e)}_{\alpha,a_{\omega_l}}(t_{l-1} - t_{e-1}) e^{-\rho t_{l-1}},\end{aligned} \tag{15}$$

with $e = \epsilon(t_{l-1} \wedge t_l)$. The second term in (14) represents the case when recombination decouples the lineages at the two loci, and they are both absorbed independently. It yields

$$\begin{aligned}
&\int_{t_b=0}^{t_{l-1}\wedge t_l} \mathbb{P}\{T_{l-1}^A \in dt_{l-1}, G_{l-1} = \omega_{l-1}, X_{l-1} = x_{l-1}, T_l^A \in dt_l, G_l = \omega_l, X_l = x_l, T^B \in dt_b\} \\
&\quad = \frac{1}{n_{\omega_{l-1}}} \frac{1}{n_{\omega_l}} \\
&\quad\quad \times \Bigg( \sum_{\epsilon=1}^{e-1} \int_{t_b=t_{\epsilon-1}}^{t_\epsilon} \sum_{\eta \in \Gamma_\epsilon} \mathbb{P}\{T_{l-1}^A \in dt_{l-1}, G_{l-1} = \omega_{l-1}, T_l^A \in dt_l, G_l = \omega_l, T^B \in dt_b, G_{t_b}^B = \eta\} \\
&\quad\quad\quad + \int_{t_b=t_{e-1}}^{t_{l-1}\wedge t_l} \sum_{\eta \in \Gamma_\epsilon} \mathbb{P}\{T_{l-1}^A \in dt_{l-1}, G_{l-1} = \omega_{l-1}, T_l^A \in dt_l, G_l = \omega_l, T^B \in dt_b, G_{t_b}^B = \eta\} \Bigg),
\end{aligned} \tag{16}$$

which is partitioned with respect to the epoch $\epsilon$ during which the recombination event occurred and the random sub-population $\{G_{t_b}^B = \eta\}$ that the coupled additional lineages were residing in at the time of the event. Note that in this partitioning, only the epochs of positive length have to be considered, since the probability of recombination in an epoch of length zero is zero.

For a given epoch $\epsilon$ of positive length, partitioning with respect to the possible sub-populations at the beginning and the end of epoch $\epsilon$, the inner term of the first summand in (16) yields

$$\begin{aligned}
&\int_{t_b=t_{\epsilon-1}}^{t_\epsilon} \sum_{\eta \in \Gamma_\epsilon} \mathbb{P}\{T_{l-1}^A \in dt_{l-1}, G_{l-1} = \omega_{l-1}, T_l^A \in dt_l, G_l = \omega_l, T^B \in dt_b, G_{t_b}^B = \eta\} \\
&= \int_{t_b=t_{\epsilon-1}}^{t_\epsilon} \mathbb{P}\{T^B \in dt_b\} \sum_{\eta \in \Gamma_\epsilon} \mathbb{P}\{T_{l-1}^A \in dt_{l-1}, G_{l-1} = \omega_{l-1}, T_l^A \in dt_l, G_l = \omega_l, G_{t_b}^B = \eta | T^B \in dt_b\} \\
&= \int_{t_b=t_{\epsilon-1}}^{t_\epsilon} \rho e^{-\rho t_b} \sum_{\eta \in \Gamma_\epsilon} \sum_{\gamma_\epsilon \in \Gamma_\epsilon} p_{\alpha,\gamma_\epsilon}^{(0,\epsilon-1)} \big(e^{(t_b - t_{\epsilon-1})Z_\epsilon}\big)_{\gamma_\epsilon,\eta} \\
&\qquad\qquad \times \sum_{Z_{\epsilon+1} \in \Gamma_{\epsilon+1}} \sum_{\substack{\zeta_\epsilon \in \Gamma_\epsilon \\ \zeta_\epsilon \subset Z_{\epsilon+1}}} \big(e^{(t_\epsilon - t_b)Z_\epsilon}\big)_{\eta,\zeta_\epsilon} q_{Z_{\epsilon+1}, a_{\omega_{l-1}}}^{(\epsilon,\epsilon(t_{l-1}))} (t_{l-1} - t_{\epsilon(t_{l-1})-1}) \\
&\qquad\qquad \times \sum_{\Xi_{\epsilon+1} \in \Gamma_{\epsilon+1}} \sum_{\substack{\xi_\epsilon \in \Gamma_\epsilon \\ \xi_\epsilon \subset \Xi_{\epsilon+1}}} \big(e^{(t_\epsilon - t_b)Z_\epsilon}\big)_{\eta,\xi_\epsilon} q_{\Xi_{\epsilon+1}, a_{\omega_l}}^{(\epsilon,\epsilon(t_l))} (t_l - t_{\epsilon(t_l)-1}) \, dt_b \\
&= \sum_{\gamma_\epsilon \in \Gamma_\epsilon} \sum_{Z_{\epsilon+1} \in \Gamma_{\epsilon+1}} \sum_{\substack{\zeta_\epsilon \in \Gamma_\epsilon \\ \zeta_\epsilon \subset Z_{\epsilon+1}}} \sum_{\Xi_{\epsilon+1} \in \Gamma_{\epsilon+1}} \sum_{\substack{\xi_\epsilon \in \Gamma_\epsilon \\ \xi_\epsilon \subset \Xi_{\epsilon+1}}} p_{\alpha,\gamma_\epsilon}^{(0,\epsilon-1)} \\
&\qquad \times R_{\gamma_\epsilon,(\zeta_\epsilon,\xi_\epsilon)}^{(\epsilon)} q_{Z_{\epsilon+1}, a_{\omega_{l-1}}}^{(\epsilon,\epsilon(t_{l-1}))} (t_{l-1} - t_{\epsilon(t_{l-1})-1}) q_{\Xi_{\epsilon+1}, a_{\omega_l}}^{(\epsilon,\epsilon(t_l))} (t_l - t_{\epsilon(t_l)-1}),
\end{aligned} \tag{17}$$

where

$$\begin{aligned}
R_{\gamma_\epsilon,(\zeta_\epsilon,\xi_\epsilon)}^{(\epsilon)} &:= \sum_{\eta \in \Gamma_\epsilon} \int_{t_b=t_{\epsilon-1}}^{t_\epsilon} \rho e^{-\rho t_b} \big(e^{(t_b - t_{\epsilon-1})Z_\epsilon}\big)_{\gamma_\epsilon,\eta} \big(e^{(t_\epsilon - t_b)Z_\epsilon}\big)_{\eta,\zeta_\epsilon} \big(e^{(t_\epsilon - t_b)Z_\epsilon}\big)_{\eta,\xi_\epsilon} \, dt_b \\
&= \rho \sum_{\eta \in \Gamma_\epsilon} \sum_{k=1}^{2g_\epsilon} \sum_{m=1}^{2g_\epsilon} \sum_{n=1}^{2g_\epsilon} (v_k^{(\epsilon)} w_k^{(\epsilon)})_{\gamma_\epsilon,\eta} (v_m^{(\epsilon)} w_m^{(\epsilon)})_{\eta,\zeta_\epsilon} (v_n^{(\epsilon)} w_n^{(\epsilon)})_{\eta,\xi_\epsilon} \\
&\qquad \times \int_{t_b=t_{\epsilon-1}}^{t_\epsilon} e^{-\rho t_b} e^{\lambda_k^{(\epsilon)}(t_b - t_{\epsilon-1})} e^{\lambda_m^{(\epsilon)}(t_\epsilon - t_b)} e^{\lambda_n^{(\epsilon)}(t_\epsilon - t_b)} \, dt_b \\
&= \rho \sum_{\eta \in \Gamma_\epsilon} \sum_{k=1}^{2g_\epsilon} \sum_{m=1}^{2g_\epsilon} \sum_{n=1}^{2g_\epsilon} (v_k^{(\epsilon)} w_k^{(\epsilon)})_{\gamma_\epsilon,\eta} (v_m^{(\epsilon)} w_m^{(\epsilon)})_{\eta,\zeta_\epsilon} (v_n^{(\epsilon)} w_n^{(\epsilon)})_{\eta,\xi_\epsilon} \\
&\qquad \times H_{t_{\epsilon-1}}^{t_\epsilon}((\lambda_m^{(\epsilon)} + \lambda_n^{(\epsilon)}) t_\epsilon - \lambda_k^{(\epsilon)} t_{\epsilon-1}, \lambda_k^{(\epsilon)} - \lambda_m^{(\epsilon)} - \lambda_n^{(\epsilon)} - \rho),
\end{aligned} \tag{18}$$

using the spectral decomposition to simplify the matrix exponentials. Note that this quantity is independent of $t_{l-1}$ and $t_l$. The definition

$$H_a^b(u,\lambda) = \int_{t=a}^{b} e^{\lambda t+u}\,dt = \begin{cases} \frac{1}{\lambda}(e^{\lambda b+u} - e^{\lambda a+u}), & \text{if } \Re(u) \neq \pm\infty,\ b \neq \infty,\ \lambda \in \mathbb{C}\backslash\{0\}, \\ e^u(b-a), & \text{if } \Re(u) \neq \pm\infty,\ b \neq \infty,\ \lambda = 0, \\ -\frac{1}{\lambda}e^{\lambda a+u}, & \text{if } \Re(u) \neq \pm\infty,\ b = \infty,\ \Re(\lambda) < 0, \\ 0, & \text{if } \Re(u) = -\infty, \end{cases} \quad [19]$$

is used for the integral term in (18), with $b > a \geq 0$. This definition covers all the relevant cases, since $\Re(\lambda_n^{(\epsilon)}) \leq 0$ holds for all $\epsilon$ and $n$, and $\Re(\lambda_n^{(\epsilon)}) = 0$ implies $\lambda_n^{(\epsilon)} = 0$ for $Z_\epsilon$ considered here. Also, whenever $\Re(u) \neq \pm\infty$ and $b = \infty$, then $\Re(\lambda) < 0$, if $\rho > 0$. In addition, for $a < \infty$ and $u \neq \infty$, the definition

$$0 \cdot H_a^\infty(u,0) = 0 \cdot \lim_{b\to\infty} H_a^b(u,0) = \lim_{b\to\infty} \left(0 \cdot H_a^b(u,0)\right) = \lim_{b\to\infty} 0 = 0 \quad [20]$$

has to be used in the appropriate cases.

Focusing on the second summand in (16), assume without loss of generality that $t_{l-1} < t_l$. For the case $\epsilon(t_{l-1}) \neq \epsilon(t_l)$,

$$\begin{aligned} &\int_{t_b=t_{e-1}}^{t_{l-1}} \sum_{\eta\in\Gamma_e} \mathbb{P}\{T_{l-1}^A \in dt_{l-1}, G_{l-1} = \omega_{l-1}, T_l^A \in dt_l, G_l = \omega_l, T^B \in dt_b, G_{t_b}^B = \eta\} \\ &\quad = \int_{t_b=t_{e-1}}^{t_{l-1}} \mathbb{P}\{T^B \in dt_b\} \sum_{\eta\in\Gamma_e} \mathbb{P}\{T_{l-1}^A \in dt_{l-1}, G_{l-1} = \omega_{l-1}, T_l^A \in dt_l, G_l = \omega_l, G_{t_b}^B = \eta | T^B \in dt_b\} \\ &\quad = \int_{t_b=t_{e-1}}^{t_{l-1}} \rho e^{-\rho t_b} \sum_{\eta_e\in\Gamma_e} \sum_{\gamma\in\Gamma_e} p_{\alpha,\gamma_e}^{(0,e-1)} \left(e^{(t_b-t_{e-1})Z_e}\right)_{\gamma_e,\eta} \left(Z_e e^{(t_{l-1}-t_b)Z_e}\right)_{\eta,a_{\omega_{l-1}}} \\ &\qquad\qquad \times \sum_{\Xi_{e+1}\in\Gamma_{e+1}} \sum_{\substack{\xi_e\in\Gamma_e \\ \xi_e\subset\Xi_{e+1}}} \left(e^{(t_e-t_b)Z_e}\right)_{\eta,\xi_e} q_{\Xi_{e+1},a_{\omega_l}}^{(e,\epsilon(t_l))}(t_l - t_{\epsilon(t_l)-1})\,dt_b, \end{aligned} \quad [21]$$

holds, whereas the summand yields

$$\begin{aligned} &\int_{t_b=t_{e-1}}^{t_{l-1}\wedge t_l} \rho e^{-\rho t_b} \sum_{\eta\in\Gamma_e} \sum_{\gamma_e\in\Gamma_e} p_{\alpha,\gamma_e}^{(0,e-1)} \left(e^{(t_b-t_{e-1})Z_e}\right)_{\gamma_e,\eta} \\ &\qquad \times \left(Z_e e^{(t_{l-1}-t_b)Z_e}\right)_{\eta,a_{\omega_{l-1}}} \left(Z_e e^{(t_l-t_b)Z_e}\right)_{\eta,a_{\omega_l}}\,dt_b \end{aligned} \quad [22]$$

for the case $\epsilon(t_{l-1}) = \epsilon(t_l)$. It is possible to obtain more explicit expressions for the integrals in (21) and (22), and to compute them numerically, using the spectral decompositions introduced in Section 1.3.3. However, we will not provide these computations here, but rather provide the full details for the discretized HMM underlying $\pi_\Theta^D$ in the next section. Finally, note that we provided the details for computing the joint density for the absorbing lineages at locus $l-1$ and $l$ here, but dividing this density by the marginal density at locus $l-1$ yields the requisite transition density for the HMM.

***1.4.3. Emission.*** Conditional on the absorption time $t_l$, the number of mutation events is Poisson distributed with parameter $\theta$, the mutation rate. Thus, the emission probability is given as

$$\begin{aligned} &\mathbb{P}\{H[l] = a | T_l^A \in dt_l, G_l = \omega_l, X_l = x_l\} \\ &\quad = \left(e^{t_l\theta(P-\mathbb{1})}\right)_{x_l[l],a}, \end{aligned} \quad [23]$$

where $H$ denotes the additionally sampled haplotype, $a \in E$, $x_l[l]$ is the allele that the absorbing lineage bears at locus $l$, and $P$ is the mutation matrix that governs the transitions between the alleles.

**1.5. Discretized HMM.** To compute the approximate CSD $\pi_\Theta^T(h|\alpha, \mathbf{n})$ under the continuous model, one would have to integrate the probability of observing $h$ given a certain sequence of marginal additional genealogies over all possible such sequences. Since the HMM over this infinite state space cannot be implemented efficiently, we will introduce another approximation by discretizing the hidden state space. In this section, we will provide details about the initial, transition and emission probabilities for the discrete HMM underlying the CSD $\pi_\Theta^D$. The basic idea is to integrate the respective densities introduced in the previous section over the discretization intervals. For ease of notation, we will use the partition of the past into demographic epochs as the discretization of the absorption time. However, we describe in Section 6 how this restriction can be relaxed. Furthermore, note that the epochs of length zero do not yield valid hidden states of the discretized HMM.

The hidden state space then comprises of an epoch of absorption $i \in \mathcal{E}$, a population where absorption takes place $\omega \in \Gamma_i$, and an absorbing trunk-lineage $x \in \mathbf{n}_\omega$. For arbitrary hidden states $s_l = (i_l, \omega_l, x_l)$ and $s_{l-1} = (i_{l-1}, \omega_{l-1}, x_{l-1})$, the initial probabilities

$$\nu(s_l) := \mathbb{P}\{T_l^A \in I_{i_l}, G_l = \omega_l, X_l = x_l\}, \qquad [24]$$

the transition probabilities

$$\begin{aligned}\phi(s_l|s_{l-1}) := \mathbb{P}\{&T_l^A \in I_{i_l}, G_l = \omega_l, X_l = x_l | \\ &T_{l-1}^A \in I_{i_{l-1}}, G_{l-1} = \omega_{l-1}, X_{l-1} = x_{l-1}\}\end{aligned} \qquad [25]$$

and the emission probabilities

$$\xi(h[l]|s_l) := \mathbb{P}\{H[l] = h[l] | T_l^A \in I_{i_l}, G_l = \omega_l, X_l = x_l\}, \qquad [26]$$

can be computed using suitable combinations of the matrix exponentials describing the evolution of the Markov chain that governs the dynamics of the marginal additional lineage backwards in time. In the following sections, we provide the details of these computations. Note that the requisite Markov chain becomes inhomogeneous when considering more general population size models such as exponential growth. The solution then cannot be obtained by matrix exponentials, but we rather have to resort to numerical approximations via step-wise solving of the associated differential equations. Both procedures are implemented in our software package. A realization of the discretized CSD can be seen in Figure S6B.

***1.5.1. Marginal/Initial probability.*** The probability that the additional lineage residing in sub-population $\Xi$ at time $t_\epsilon$ is absorbed into any lineage of the trunk within the sub-population $\omega$ during the interval $I_i$ is given by

$$\begin{aligned}Q_{\Xi,a_\omega}^{(\epsilon)}(i) &:= \int_{t=t_{i-1}}^{t_i} q_{\Xi,a_\omega}^{(\epsilon,i)}(t - t_{i-1})\, dt \\ &= \int_{t=t_{i-1}}^{t_i} \sum_{\gamma_i \in \Gamma_i} p_{\Xi,\gamma_i}^{(\epsilon,i-1)} \big(Z_i e^{(t-t_{i-1})Z_i}\big)_{\gamma_i,a_\omega}\, dt \\ &= \sum_{\gamma_i \in \Gamma_i} p_{\Xi,\gamma_i}^{(\epsilon,i-1)} \sum_{k=1}^{2g_i} (v_k^{(i)} w_k^{(i)})_{\gamma_i,a_\omega} \lambda_k^{(i)} e^{-\lambda_k^{(i)} t_{i-1}} \int_{t=t_{i-1}}^{t_i} e^{\lambda_k^{(i)} t}\, dt \\ &= \sum_{\gamma_i \in \Gamma_i} p_{\Xi,\gamma_i}^{(\epsilon,i-1)} \sum_{k=1}^{2g_i} (v_k^{(i)} w_k^{(i)})_{\gamma,a_\omega} \lambda_k^{(i)} e^{-\lambda_k^{(i)} t_{i-1}} H_{t_{i-1}}^{t_i}(0, \lambda_k^{(i)})\end{aligned} \qquad [27]$$

where we used $q$ as defined in (13), and $H$ as in (19) and (20). The discretized initial probability of being absorbed during the interval $I_i$ in sub-population $\omega_l \in \Gamma_i$ into the lineage $x_l \in \mathbf{n}_{\omega_l}$ is then given as

$$\nu(\omega_l, i, x_l) := \mathbb{P}\{T_l^A \in I_i, G_l = \omega_l, X_l = x_l\} = \frac{1}{n_{\omega_l}} u(\omega_l, i) \qquad [28]$$

with

$$u(\omega, i) := \int_{t=t_{i-1}}^{t_i} q^{(0,i)}_{\alpha,a_\omega}(t - t_{i-1})\, dt = Q^{(0)}_{\alpha,a_\omega}(i). \qquad [29]$$

***1.5.2. Transition probability.*** To derive the transition probability, for given $i, j \in \mathcal{E}$, we again start by focusing on the joint probability that the additional lineage at locus $l-1$ is absorbed during the interval $I_i$ into the trunk-lineage $x_{l-1} \in \mathbf{n}_{\omega_{l-1}}$ residing in sub-population $\omega_{l-1} \in \Gamma_i$, and the lineage at locus $l$ is absorbed during $I_j$ into $x_l \in \mathbf{n}_{\omega_l}$, with $\omega_l \in \Gamma_j$. This probability is given by

$$\begin{aligned}
&\mathbb{P}\{T^A_{l-1} \in I_i, G_{l-1} = \omega_{l-1}, X_{l-1} = x_{l-1}, T^A_l \in I_j, G_l = \omega_l, X_l = x_l\} \\
&\quad = \int_{t_{l-1}=t_{i-1}}^{t_i} \int_{t_l=t_{j-1}}^{t_j} \mathbb{P}\{T^A_{l-1} \in dt_{l-1}, G_{l-1} = \omega_{l-1}, X_{l-1} = x_{l-1}, T^A_l \in dt_l, G_l = \omega_l, X_l = x_l\}.
\end{aligned} \qquad [30]$$

Substituting (14) into (30), the *coupled* term isolated in (15) yields

$$\begin{aligned}
&\int_{t_{l-1}=t_{i-1}}^{t_i} \int_{t_l=t_{j-1}}^{t_j} \delta(t_{l-1} - t_l)\delta_{\omega_{l-1},\omega_l}\delta_{x_{l-1},x_l} e^{-\rho t_{l-1}} q^{(0,\mu)}_{\alpha,a_{\omega_{l-1}}}(t_{l-1} - t_{\mu-1}) \frac{1}{n_{\omega_{l-1}}}\, dt_{l-1}\, dt_l \\
&\quad = \int_{t_{l-1}=t_{i-1}}^{t_i} \mathbb{1}_{\{t_{l-1}\in I_j\}} \delta_{\omega_{l-1},\omega_l}\delta_{x_{l-1},x_l} e^{-\rho t_{l-1}} q^{(0,\mu)}_{\alpha,a_{\omega_l}}(t_{l-1} - t_{\mu-1}) \frac{1}{n_{\omega_{l-1}}}\, dt_{l-1} \\
&\quad = \delta_{i,j}\delta_{\omega_{l-1},\omega_l}\delta_{x_{l-1},x_l} \frac{1}{n_{\omega_{l-1}}} \int_{t_{l-1}=t_{i-1}}^{t_i} e^{-\rho t_{l-1}} \sum_{\gamma_\mu\in\Gamma_\mu} p^{(0,\mu-1)}_{\alpha,\gamma_\mu} \left(Z_\mu e^{(t_{l-1}-t_{\mu-1})Z_\mu}\right)_{\gamma_\mu,a_{\omega_{l-1}}} dt_{l-1} \\
&\quad = \delta_{i,j}\delta_{\omega_{l-1},\omega_l}\delta_{x_{l-1},x_l} \frac{1}{n_{\omega_{l-1}}} \sum_{\gamma_\mu\in\Gamma_\mu} p^{(0,\mu-1)}_{\alpha,\gamma_\mu} \\
&\qquad\qquad \times \sum_{k=1}^{2g_\mu} (v^{(\mu)}_k w^{(\mu)}_k)_{\gamma_\mu,a_{\omega_{l-1}}} \lambda^{(\mu)}_k e^{-\lambda^{(\mu)}_k t_{\mu-1}} \int_{t_{l-1}=t_{i-1}}^{t_i} e^{-\rho t_{l-1}} e^{\lambda^{(\mu)}_k t_{l-1}}\, dt_{l-1} \\
&\quad = \delta_{i,j}\delta_{\omega_{l-1},\omega_l}\delta_{x_{l-1},x_l} \frac{1}{n_{\omega_{l-1}}} \sum_{\gamma_\mu\in\Gamma_\mu} p^{(0,\mu-1)}_{\alpha,\gamma_\mu} \\
&\qquad\qquad \times \sum_{k=1}^{2g_\mu} (v^{(\mu)}_k w^{(\mu)}_k)_{\gamma_\mu,a_{\omega_{l-1}}} \lambda^{(\mu)}_k H^{t_\mu}_{t_{\mu-1}}(-\lambda^{(\mu)}_k t_{\mu-1}, \lambda^{(\mu)}_k - \rho),
\end{aligned} \qquad [31]$$

with $\mu = i \wedge j$, and $H$ as defined in (19) and (20).

Furthermore, after substituting, the term in parentheses in the *decoupled* term isolated in (16) yields

$$\begin{aligned}
&\int_{t_{l-1}=t_{i-1}}^{t_i} \int_{t_l=t_{j-1}}^{t_j} \Bigg( \sum_{\epsilon=1}^{\mu-1} \int_{t_b=t_{\epsilon-1}}^{t_\epsilon} \sum_{\eta\in\Gamma_\epsilon} \mathbb{P}\{T^A_{l-1} \in dt_{l-1}, G_{l-1} = \omega_{l-1}, T^A_l \in dt_l, G_l = \omega_l, T^B \in dt_b, G^B_{t_b} = \eta\} \\
&\qquad\qquad + \int_{t_b=t_{\mu-1}}^{t_{l-1}\wedge t_l} \sum_{\eta\in\Gamma_\mu} \mathbb{P}\{T^A_{l-1} \in dt_{l-1}, G_{l-1} = \omega_{l-1}, T^A_l \in dt_l, G_l = \omega_l, T^B \in dt_b, G^B_{t_b} = \eta\} \Bigg).
\end{aligned} \qquad [32]$$

Again, fixing $\epsilon$ and focusing on a summand of the first sum in (32) yields

$$\begin{aligned}
&\int_{t_b=t_{\epsilon-1}}^{t_\epsilon} \rho e^{-\rho t_b} \sum_{\eta\in\Gamma_\epsilon}\sum_{\gamma_\epsilon\in\Gamma_\epsilon} p^{(0,\epsilon-1)}_{\alpha,\gamma_\epsilon}\big(e^{(t_b-t_{\epsilon-1})Z_\epsilon}\big)_{\gamma_\epsilon,\eta} \\
&\quad\times \sum_{Z_{\epsilon+1}\in\Gamma_{\epsilon+1}} \sum_{\substack{\zeta_\epsilon\in\Gamma_\epsilon\\ \zeta_\epsilon\subset Z_{\epsilon+1}}} \big(e^{(t_\epsilon-t_b)Z_\epsilon}\big)_{\eta,\zeta_\epsilon} \int_{t_{l-1}=t_{i-1}}^{t_i} q^{(\epsilon,i)}_{Z_{\epsilon+1},a_{\omega_{l-1}}}(t_{l-1}-t_{i-1})\,dt_{l-1} \\
&\quad\times \sum_{\Xi_{\epsilon+1}\in\Gamma_{\epsilon+1}} \sum_{\substack{\xi_\epsilon\in\Gamma_\epsilon\\ \xi_\epsilon\subset \Xi_{\epsilon+1}}} \big(e^{(t_\epsilon-t_b)Z_\epsilon}\big)_{\eta,\xi_\epsilon} \int_{t_l=t_{j-1}}^{t_j} q^{(\epsilon,j)}_{\Xi_{\epsilon+1},a_{\omega_l}}(t_l-t_{j-1})\,dt_l\,dt_b \\
&= \sum_{\gamma_\epsilon\in\Gamma_\epsilon}\sum_{Z_{\epsilon+1}\in\Gamma_{\epsilon+1}}\sum_{\Xi_{\epsilon+1}\in\Gamma_{\epsilon+1}} p^{(0,\epsilon-1)}_{\alpha,\gamma_\epsilon} Q^{(\epsilon)}_{Z_{\epsilon+1},a_{\omega_{l-1}}}(i) Q^{(\epsilon)}_{\Xi_{\epsilon+1},a_{\omega_l}}(j) \\
&\quad\times \sum_{\eta\in\Gamma_\epsilon}\sum_{\substack{\zeta_\epsilon\in\Gamma_\epsilon\\ \zeta_\epsilon\subset Z_{\epsilon+1}}}\sum_{\substack{\xi_\epsilon\in\Gamma_\epsilon\\ \xi_\epsilon\subset \Xi_{\epsilon+1}}} \int_{t_b=t_{\epsilon-1}}^{t_\epsilon} \rho e^{-\rho t_b}\big(e^{(t_b-t_{\epsilon-1})Z_\epsilon}\big)_{\gamma_\epsilon,\eta}\big(e^{(t_\epsilon-t_b)Z_\epsilon}\big)_{\eta,\zeta_\epsilon}\big(e^{(t_\epsilon-t_b)Z_\epsilon}\big)_{\eta,\xi_\epsilon}\,dt_b \\
&= \sum_{\gamma_\epsilon\in\Gamma_\epsilon}\sum_{Z_{\epsilon+1}\in\Gamma_{\epsilon+1}}\sum_{\Xi_{\epsilon+1}\in\Gamma_{\epsilon+1}} p^{(0,\epsilon-1)}_{\alpha,\gamma_\epsilon} R^{(\epsilon)}_{\gamma_\epsilon,(Z_{\epsilon+1},\Xi_{\epsilon+1})} Q^{(\epsilon)}_{Z_{\epsilon+1},a_{\omega_{l-1}}}(i) Q^{(\epsilon)}_{\Xi_{\epsilon+1},a_{\omega_l}}(j),
\end{aligned} \qquad [33]$$

where $Q^{(\cdot)}_{\cdot,\cdot}(\cdot)$ was defined in (27), and $R^{(\cdot)}_{\cdot,(\cdot,\cdot)}$ was defined in (18).

For the second summand in (32) the cases $i \neq j$ and $i = j$ have to be distinguished Assume without loss of generality $i < j$, so $\mu = i$. Then, focusing on the case $i < j$ gives

$$\begin{aligned}
&\int_{t_{l-1}=t_{\mu-1}}^{t_\mu}\int_{t_l=t_{j-1}}^{t_j}\int_{t_b=t_{\mu-1}}^{t_{l-1}} \rho e^{-\rho t_b}\sum_{\eta\in\Gamma_\mu}\sum_{\gamma_\mu\in\Gamma_\mu} p^{(0,\mu-1)}_{\alpha,\gamma_\mu}\big(e^{(t_b-t_{\mu-1})Z_\mu}\big)_{\gamma_\mu,\eta}\big(Z_\mu e^{(t_{l-1}-t_b)Z_\mu}\big)_{\eta,a_{\omega_{l-1}}} \\
&\times \sum_{\Xi_{\mu+1}\in\Gamma_{\mu+1}}\sum_{\substack{\xi_\mu\in\Gamma_\mu\\ \xi_\mu\subset\Xi_{\mu+1}}} \big(e^{(t_\mu-t_b)Z_\mu}\big)_{\eta,\xi_\mu} q^{(\mu,j)}_{\Xi_{\mu+1},a_{\omega_l}}(t_l-t_{j-1})\,dt_b\,dt_l\,dt_{l-1} \\
&= \sum_{\eta\in\Gamma_\mu}\sum_{\gamma_\mu\in\Gamma_\mu}\sum_{\Xi_{\mu+1}\in\Gamma_{\mu+1}}\sum_{\substack{\xi_\mu\in\Gamma_\mu\\ \xi_\mu\subset\Xi_{\mu+1}}} p^{(0,\mu-1)}_{\alpha,\gamma_\mu} Q^{(\mu)}_{\Xi_{\mu+1},a_{\omega_l}}(j) \\
&\times \int_{t_{l-1}=t_{\mu-1}}^{t_\mu}\int_{t_b=t_{\mu-1}}^{t_{l-1}} \rho e^{-\rho t_b}\big(e^{(t_b-t_{\mu-1})Z_\mu}\big)_{\gamma_\mu,\eta}\big(e^{(t_\mu-t_b)Z_\mu}\big)_{\eta,\xi_\mu}\big(Z_\mu e^{(t_{l-1}-t_b)Z_\mu}\big)_{\eta,a_{\omega_{l-1}}}\,dt_b\,dt_{l-1} \\
&= \sum_{\eta\in\Gamma_\mu}\sum_{\gamma_\mu\in\Gamma_\mu}\sum_{\Xi_{\mu+1}\in\Gamma_{\mu+1}}\sum_{\substack{\xi_\mu\in\Gamma_\mu\\ \xi_\mu\subset\Xi_{\mu+1}}} p^{(0,\mu-1)}_{\alpha,\gamma_\mu} Q^{(\mu)}_{\Xi_{\mu+1},a_{\omega_l}}(j) \\
&\times \sum_{k=1}^{2g_\mu}\sum_{m=1}^{2g_\mu}\sum_{n=1}^{2g_\mu} (v^{(\mu)}_k w^{(\mu)}_k)_{\gamma_\mu,\eta}(v^{(\mu)}_m w^{(\mu)}_m)_{\eta,\xi_\mu}(v^{(\mu)}_n w^{(\mu)}_n)_{\eta,a_{\omega_{l-1}}} \\
&\qquad\times \rho\lambda^{(\mu)}_n e^{\lambda^{(\mu)}_m t_\mu - \lambda^{(\mu)}_k t_{\mu-1}} \int_{t_{l-1}=t_{\mu-1}}^{t_\mu} e^{\lambda^{(\mu)}_n t_{l-1}} \int_{t_b=t_{\mu-1}}^{t_{l-1}} e^{(\lambda^{(\mu)}_k-\lambda^{(\mu)}_m-\lambda^{(\mu)}_n-\rho)t_b}\,dt_b\,dt_{l-1}.
\end{aligned} \qquad [34]$$

Changing the order of integration in the integral expression in (34) yields

$$\begin{aligned}
\lambda^{(\mu)}_n &\int_{t_b=t_{\mu-1}}^{t_\mu} e^{(\lambda^{(\mu)}_k-\lambda^{(\mu)}_m-\lambda^{(\mu)}_n-\rho)t_b}\int_{t_{l-1}=t_b}^{t_\mu} e^{\lambda^{(\mu)}_n t_{l-1}}\,dt_{l-1}\,dt_b \\
&= \int_{t_b=t_{\mu-1}}^{t_\mu} e^{(\lambda^{(\mu)}_k-\lambda^{(\mu)}_m-\lambda^{(\mu)}_n-\rho)t_b}\big(e^{\lambda^{(\mu)}_n t_\mu} - e^{\lambda^{(\mu)}_n t_b}\big)\,dt_b \\
&= e^{\lambda^{(\mu)}_n t_\mu}\int_{t_b=t_{\mu-1}}^{t_\mu} e^{(\lambda^{(\mu)}_k-\lambda^{(\mu)}_m-\lambda^{(\mu)}_n-\rho)t_b}\,dt_b - \int_{t_b=t_{\mu-1}}^{t_\mu} e^{(\lambda^{(\mu)}_k-\lambda^{(\mu)}_m-\rho)t_b}\,dt_b.
\end{aligned} \qquad [35]$$

Note that the second term in (35) does not depend on $n$ anymore. Thus, when substituting it back into expression (34) this term vanishes, since $\sum_{n=1}^{2g_\mu}(v_n^{(\mu)}w_n^{(\mu)})_{\eta_b,a_{\omega_{l-1}}} = (\mathbb{1})_{\eta_b,a_{\omega_{l-1}}} = 0$. The first term in (35) can be written as

$$H_{t_{\mu-1}}^{t_\mu}(\lambda_n^{(\mu)}t_\mu, \lambda_k^{(\mu)} - \lambda_m^{(\mu)} - \lambda_n^{(\mu)} - \rho), \quad [36]$$

using the definitions (19) and (20). Note that $i < j$ and thus $t_\mu < \infty$ hold, so this quantity is well defined. Substituting it into (34) yields

$$\begin{aligned}
&\sum_{\eta\in\Gamma_\mu}\sum_{\gamma_\mu\in\Gamma_\mu}\sum_{\Xi_{\mu+1}\in\Gamma_{\mu+1}}\sum_{\substack{\xi_\mu\in\Gamma_\mu\\ \xi_\mu\subset\Xi_{\mu+1}}} p_{\alpha,\gamma_\mu}^{(0,\mu-1)}Q_{\Xi_{\mu+1},a_{\omega_l}}^{(\mu)}(j)\\
&\quad\times\sum_{k=1}^{2g_\mu}\sum_{m=1}^{2g_\mu}\sum_{n=1}^{2g_\mu}(v_k^{(\mu)}w_k^{(\mu)})_{\gamma_\mu,\eta}(v_m^{(\mu)}w_m^{(\mu)})_{\eta,\xi_\mu}(v_n^{(\mu)}w_n^{(\mu)})_{\eta,a_{\omega_{l-1}}}\\
&\qquad\times\rho H_{t_{\mu-1}}^{t_\mu}((\lambda_m^{(\mu)}+\lambda_n^{(\mu)})t_\mu-\lambda_k^{(\mu)}t_{\mu-1},\lambda_k^{(\mu)}-\lambda_m^{(\mu)}-\lambda_n^{(\mu)}-\rho)\\
&=\sum_{\gamma_\mu\in\Gamma_\mu}p_{\alpha,\gamma_\mu}^{(0,\mu-1)}\sum_{\Xi_{\mu+1}\in\Gamma_{\mu+1}}\sum_{\substack{\xi_\mu\in\Gamma_\mu\\ \xi_\mu\subset\Xi_{\mu+1}}}R_{\gamma_\mu,(a_{\omega_{l-1}},\xi_\mu)}^{(\mu)}Q_{\Xi_{\mu+1},a_{\omega_l}}^{(\mu)}(j).
\end{aligned} \quad [37]$$

In the case $i = j = \mu$, the second summand in (32) gives

$$\begin{aligned}
&\int_{t_{l-1}=t_{\mu-1}}^{t_\mu}\int_{t_l=t_{\mu-1}}^{t_\mu}\int_{t_b=t_{\mu-1}}^{t_{l-1}\wedge t_l}\rho e^{-\rho t_b}\sum_{\eta\in\Gamma_\mu}\sum_{\gamma_\mu\in\Gamma_\mu}p_{\alpha,\gamma_\mu}^{(0,\mu-1)}\\
&\quad\times\left(e^{(t_b-t_{\mu-1})Z_\mu}\right)_{\gamma_\mu,\eta_b}\left(Z_\mu e^{(t_{l-1}-t_b)Z_\mu}\right)_{\eta,a_{\omega_{l-1}}}\left(Z_\epsilon e^{(t_l-t_b)Z_\mu}\right)_{\eta,a_{\omega_l}}dt_b\,dt_l\,dt_{l-1}\\
&=\sum_{\eta\in\Gamma_\epsilon}\sum_{\gamma_\mu\in\Gamma_\epsilon}p_{\alpha,\gamma_\mu}^{(0,\mu-1)}\sum_{k=1}^{2g_\mu}\sum_{m=1}^{2g_\mu}\sum_{n=1}^{2g_\mu}(v_k^{(\mu)}w_k^{(\mu)})_{\gamma_\mu,\eta}(v_m^{(\mu)}w_m^{(\mu)})_{\eta,a_{\omega_{l-1}}}(v_n^{(\mu)}w_n^{(\mu)})_{\eta,a_{\omega_l}}\\
&\quad\times\rho\lambda_m^{(\mu)}\lambda_n^{(\mu)}e^{-\lambda_k^{(\mu)}t_{\mu-1}}\int_{t_{l-1}=t_{\mu-1}}^{t_\mu}\int_{t_l=t_{\mu-1}}^{t_\mu}\int_{t_b=t_{\mu-1}}^{t_{l-1}\wedge t_l}e^{(\lambda_k^{(\mu)}-\lambda_m^{(\mu)}-\lambda_n^{(\mu)}-\rho)t_b}e^{\lambda_m^{(\mu)}t_{l-1}}e^{\lambda_n^{(\mu)}t_l}dt_b\,dt_l\,dt_{l-1}.
\end{aligned} \quad [38]$$

Again, considering only the integral part in (32), and exchanging the order of integration yields

$$\begin{aligned}
&\lambda_m^{(\mu)}\lambda_n^{(\mu)}\int_{t_b=t_{\mu-1}}^{t_\mu}e^{(\lambda_k^{(\mu)}-\lambda_m^{(\mu)}-\lambda_n^{(\mu)}-\rho)t_b}\Big[\int_{t_{l-1}=t_b}^{t_\mu}e^{\lambda_m^{(\mu)}t_{l-1}}dt_{l-1}\Big]\Big[\int_{t_l=t_b}^{t_\mu}e^{\lambda_n^{(\mu)}t_l}dt_l\Big]dt_b.\\
&=\int_{t_b=t_{\mu-1}}^{t_\mu}e^{(\lambda_k^{(\mu)}-\lambda_m^{(\mu)}-\lambda_n^{(\mu)}-\rho)t_b}\Big[e^{\lambda_m^{(\mu)}t_\mu}-e^{\lambda_m^{(\mu)}t_b}\Big]\Big[e^{\lambda_n^{(\mu)}t_\mu}-e^{\lambda_n^{(\mu)}t_b}\Big]dt_b\\
&=\int_{t_b=t_{\mu-1}}^{t_\mu}e^{(\lambda_k^{(\mu)}-\lambda_m^{(\mu)}-\lambda_n^{(\mu)}-\rho)t_b}e^{(\lambda_m^{(\mu)}+\lambda_n^{(\mu)})t_\mu}dt_b\\
&=H_{t_{\mu-1}}^{t_\mu}((\lambda_m^{(\mu)}+\lambda_n^{(\mu)})t_\mu,\lambda_k^{(\mu)}-\lambda_m^{(\mu)}-\lambda_n^{(\mu)}-\rho),
\end{aligned} \quad [39]$$

with $H$ as defined in (19) and (20). Here the second equality holds, since upon solving the brackets, only the first summand is dependent on $k$, $m$, and $n$. The other summands then vanish due to a similar argument as was used in deriving equation (36). Substituting the right hand side of (39) into (38) gives

$$\begin{aligned}
&\sum_{\eta\in\Gamma_\epsilon}\sum_{\gamma_\mu\in\Gamma_\epsilon}p_{\alpha,\gamma_\mu}^{(0,\mu-1)}\sum_{k=1}^{2g_\mu}\sum_{m=1}^{2g_\mu}\sum_{n=1}^{2g_\mu}(v_k^{(\mu)}w_k^{(\mu)})_{\gamma_\mu,\eta}(v_m^{(\mu)}w_m^{(\mu)})_{\eta,a_{\omega_{l-1}}}(v_n^{(\mu)}w_n^{(\mu)})_{\eta,a_{\omega_l}}\\
&\quad\times\rho H_{t_{\mu-1}}^{t_\mu}((\lambda_m^{(\mu)}+\lambda_n^{(\mu)})t_\mu-\lambda_k^{(\mu)}t_{\mu-1},\lambda_k^{(\mu)}-\lambda_m^{(\mu)}-\lambda_n^{(\mu)}-\rho)\\
&=\sum_{\gamma_\mu\in\Gamma_\epsilon}p_{\alpha,\gamma_\mu}^{(0,\mu-1)}R_{\gamma_\mu,(a_{\omega_{l-1}},a_{\omega_l})}^{(\mu)}.
\end{aligned} \quad [40]$$

Combining (31), (33), (37), and (40) gives the joint absorption probability (30).

The discretized transition probability can then be obtained by dividing the joint probability through the marginal probability at locus $l-1$:

$$\begin{aligned}&\phi(i_l,\omega_l,x_l|i_{l-1},\omega_{l-1},x_{l-1})\\&\quad:=\mathbb{P}\{T_l^A\in I_{i_l},G_l=\omega_l,X_l=x_l|T_{l-1}^A\in I_{i_{l-1}},G_{l-1}=\omega_{l-1},X_{l-1}=x_{l-1}\}\\&\quad=y(i_{l-1},\omega_{l-1})\delta_{i_{l-1},i_l}\delta_{\omega_{l-1},\omega_l}\delta_{x_{l-1},x_l}+z(i_l,\omega_l|i_{l-1},\omega_{l-1})\frac{1}{n_{\omega_l}}.\end{aligned} \quad [41]$$

Here

$$y(i,\omega_{l-1}):=\frac{1}{u(i,\omega_{l-1})}\sum_{\gamma_i\in\Gamma_i}p_{\alpha,\gamma_i}^{(0,i-1)}\sum_{k=1}^{2g_i}(v_k^{(i)}w_k^{(i)})_{\gamma_i,a_{\omega_{l-1}}}\lambda_k^{(i)}H_{t_{i-1}}^{t_i}(-\lambda_k^{(i)}t_{i-1},\lambda_k^{(i)}-\rho), \quad [42]$$

with $u(\cdot,\cdot)$ as defined in (29). Furthermore, with $\mu=i\wedge j$, define

$$\begin{aligned}z(j,\omega_l|i,\omega_{l-1}):=\frac{1}{u(i,\omega_{l-1})}\Bigg[&\sum_{\epsilon=1}^{\mu-1}\sum_{\gamma_\epsilon\in\Gamma_\epsilon}p_{\alpha,\gamma_\epsilon}^{(0,\epsilon-1)}\sum_{Z_{\epsilon+1}\in\Gamma_{\epsilon+1}}\sum_{\substack{\zeta_\epsilon\in\Gamma_\epsilon\\ \zeta_\epsilon\subset Z_{\epsilon+1}}}\sum_{\Xi_{\epsilon+1}\in\Gamma_{\epsilon+1}}\sum_{\substack{\xi_\epsilon\in\Gamma_\epsilon\\ \xi_\epsilon\subset\Xi_{\epsilon+1}}}R_{\gamma_\epsilon,(\zeta_\epsilon,\xi_\epsilon)}^{(\epsilon)}\\&\times Q_{Z_{\epsilon+1},a_{\omega_{l-1}}}^{(\epsilon)}(i)Q_{\Xi_{\epsilon+1},a_{\omega_l}}^{(\epsilon)}(j)\\&+\sum_{\gamma_\mu\in\Gamma_\mu}p_{\alpha,\gamma_\mu}^{(0,\mu-1)}W_{\gamma_\mu,(a_{\omega_{l-1}},a_{\omega_l})}^{(\mu)}(i,j)\Bigg],\end{aligned} \quad [43]$$

where

$$W_{\gamma_\mu,(a_{\omega_{l-1}},a_{\omega_l})}^{(\mu)}(i,j):=\begin{cases}\sum_{\Xi_{\mu+1}\in\Gamma_{\mu+1}}\sum_{\substack{\xi_\mu\in\Gamma_\mu\\ \xi_\mu\subset\Xi_{\mu+1}}}R_{\gamma_\mu,(a_{\omega_{l-1}},\xi_\mu)}^{(\mu)}Q_{\Xi_{\mu+1},a_{\omega_l}}^{(\mu)}(j), & \text{if } i<j,\\ R_{\gamma_\mu,(a_{\omega_{l-1}},a_{\omega_l})}^{(\mu)}, & \text{if } i=j,\\ W_{\gamma_\mu,(a_{\omega_l},a_{\omega_{l-1}})}^{(\mu)}(j,i), & \text{if } i>j.\end{cases} \quad [44]$$

***1.5.3. Emission probability.*** Finally, the emission probability, that is, the probability that the observed haplotype $H$ carries the allele $a$ at locus $l$ given that the additional lineage at this locus is absorbed during the interval $I_i$ in sub-population $\omega_l\in\Gamma_i$ into the lineage $x_l\in\mathbf{n}_{\omega_l}$ can be computed as

$$\begin{aligned}&\mathbb{P}\{H[l]=a|T_l^A\in I_i,G_l=\omega_l,X_l=x_l\}\\&=\frac{\mathbb{P}\{H[l]=a,T_l^A\in I_i,G_l=\omega_l,X_l=x_l\}}{\mathbb{P}\{T_l^A\in I_i,G_l=\omega_l,X_l=x_l\}}\\&=\frac{\mathbb{P}\{H[l]=a,T_l^A\in I_i,G_l=\omega_l,X_l=x_l\}}{u(i,\omega_l)}n_{\omega_l}.\end{aligned} \quad [45]$$

Using (23) and (13), the numerator in (45) yields

$$\begin{aligned}&\mathbb{P}\{H[l]=a,T_l^A\in I_i,G_l=\omega_l,X_l=x_l\}n_{\omega_l}\\&\quad=\int_{t_l=t_{i-1}}^{t_i}\mathbb{P}\{H[l]=a,T_l^A\in dt_l,G_l=\omega_l,X_l=x_l\}n_{\omega_l}\\&\quad=\int_{t_l=t_{i-1}}^{t_i}\mathbb{P}\{H[l]=a|T_l^A\in dt_l,G_l=\omega_l,X_l=x_l\}\mathbb{P}\{T_l^A\in dt_l,G_l=\omega_l,X_l=x_l\}n_{\omega_l}\\&\quad=\int_{t_l=t_{i-1}}^{t_i}(e^{t_l\theta(P-\mathbb{1})})_{x_l[l],a}\sum_{\gamma_i\in\Gamma_i}p_{\alpha,\gamma_i}^{(0,i-1)}(Z_ie^{(t_l-t_{i-1})Z_i})_{\gamma_i,a_{\omega_l}}\,dt_l\\&\quad=\sum_{\gamma_i\in\Gamma_i}p_{\alpha,\gamma_i}^{(0,i-1)}\sum_{j=1}^{|E|}\sum_{k=1}^{2g_i}(\mathfrak{v}_j\mathfrak{w}_j)_{x_l[l],a}(v_k^{(i)}w_k^{(i)})_{\gamma_i,a_{\omega_l}}\lambda_k^{(i)}H_{t_{i-1}}^{t_i}\big(-\lambda_k^{(i)}t_{i-1},\theta(\mathfrak{l}_j-1)+\lambda_k^{(i)}\big),\end{aligned} \quad [46]$$

where $\mathfrak{l}_j$ (with $\Re(\mathfrak{l}_j) \leq 0$) are the eigenvalues of $P$, $\mathfrak{v}_j$ are its eigenvectors, and $\mathfrak{w}_j$ the are row-vectors of the matrix inverse to the matrix made up of the column vectors $\mathfrak{v}_j$. Again, $H$ is as defined in (19) and (20). Combining (45) with (46) yields, with $u(\cdot,\cdot)$ as defined in (29),

$$\begin{aligned}\xi(a|i,\omega_l,x_l) &:= \mathbb{P}\{H[l]=a|T_l^A \in I_i, G_l=\omega_l, X_l=x_l\}\\ &= \frac{1}{u(i,\omega_l)} \sum_{\gamma_i\in\Gamma_i} p^{(0,i-1)}_{\alpha,\gamma_i} \sum_{j=1}^{|E|}\sum_{k=1}^{2g_i} (\mathfrak{v}_j\mathfrak{w}_j)_{x_l[l],a} (v_k^{(i)} w_k^{(i)})_{\gamma_i,a_{\omega_l}} \lambda_k^{(i)} \\ &\qquad \times H_{t_{i-1}}^{t_i}\big(-\lambda_k^{(i)} t_{i-1}, \theta(\mathfrak{l}_j-1)+\lambda_k^{(i)}\big)\end{aligned} \quad [47]$$

for the emission probability of the discretized HMM underlying the CSD $\pi_\Theta^D$.

### 2. Forward-Backward Algorithm

Given a certain demographic history $\Theta$ and an observed configuration $\mathbf{n}$, denote by $H_\Theta^{\alpha,\mathbf{n}} \in E^L$ the random haplotype additionally sampled in sub-population $\alpha$ which is distributed according to the CSD $\pi_\Theta^D$, that is, $H_\Theta^{\alpha,\mathbf{n}} \sim \pi_\Theta^D(\cdot|\alpha,\mathbf{n})$. Note that the distribution implicitly depends on the recombination rate $\rho$ and the mutational model $(\theta, P)$ as well. The probability $\mathbb{P}\{H_\Theta^{\alpha,\mathbf{n}} = h\}$ of observing a certain additional haplotype $h \in E^L$ can be computed under the HMM defined by the probabilities $\nu$, $\phi$, and $\xi$ given in Section 1.5 using the forward algorithm. To this end denote by $\mathcal{S} := \big\{(i,\omega,x)\big| i\in\mathcal{E}, \omega\in\Gamma_i, x\in\mathbf{n}_\omega\big\}$ the set of hidden states, so a hidden state comprises of an interval $i$ during which the additional lineage is absorbed, a sub-population $\omega$ in which absorption happens, and a trunk-lineage $x$ that the lineage is absorbed into. Further, for $1 \leq l \leq L$, denote by $S_l \in \mathcal{S}$ the random hidden state at locus $l$, and by $S_\Theta^{\alpha,\mathbf{n}} := (S_1,\ldots,S_L)$ the full sequence of hidden states.

#### 2.1. Forward Algorithm.

Given the hidden state $s_l = (i_l,\omega_l,x_l) \in \mathcal{S}$, the forward probability

$$\begin{aligned}F_l(s_l) &:= \mathbb{P}\{H_\Theta^{\alpha,\mathbf{n}}[1:l] = h[1:l], S_l = s_l\}\\ &= \mathbb{P}\{H_\Theta^{\alpha,\mathbf{n}}[1:l] = h[1:l], T_l^A \in I_{i_l}, G_l=\omega_l, X_l=x_l\}\end{aligned} \quad [48]$$

is the joint probability of observing the partial haplotype $h[1:l]$ up to locus $l$, and the additional lineage being absorbed into haplotype $x_l$ in sub-population $\omega_l$ during interval $i_l$ at locus $l$. Dynamic programming can be used to compute $F_l(s_l)$ via the dynamic program:

$$\begin{aligned}F_l(s_l) &= \xi(h_l|s_l) \sum_{s_{l-1}\in\mathcal{S}} F_{l-1}(s_{l-1})\phi(s_l|s_{l-1})\\ &= \xi(h_l|i_l,\omega_l,x_l)\Big[y(i_l,\omega_l)F_{l-1}(i_l,\omega_l,x_l)\\ &\qquad + \frac{1}{n_{\omega_l}} \sum_{\substack{i_{l-1}\in\mathcal{E},\\ \omega_{l-1}\in\Gamma_{i_{l-1}}}} z(i_l,\omega_l|i_{l-1},\omega_{l-1}) \sum_{x_{l-1}\in\mathbf{n}_{\omega_{l-1}}} F_{l-1}(i_{l-1},\omega_{l-1},x_{l-1})\Big].\end{aligned} \quad [49]$$

The initial value for this dynamic program is given by

$$F_1(i_1,\omega_1,x_1) = \xi_\theta(h_1|i_1,\omega_1,x_1,)\frac{1}{n_{\omega_1}}u(i_1,\omega_1). \quad [50]$$

Note that if the haplotypes associated with lineages $x$ and $x'$ from the trunk are identical, then we have $F_l(i,\omega,x) = F_l(i,\omega,x')$ for all $l,i,\omega$. Finally, the probability of observing the additional haplotype is given as

$$\mathbb{P}\{H_\Theta^{\alpha,\mathbf{n}} = h\} = \sum_{s_L\in\mathcal{S}} \mathbb{P}\{H_\Theta^{\alpha,\mathbf{n}}[1:L] = h_{[1:L]}, S_L = s_L\} = \sum_{s_L\in\mathcal{S}} F_L(s_L). \quad [51]$$

A naïve implementation of (49) would, for all $s_l \in \mathcal{S}$, iterate over every $s_{l-1} \in \mathcal{S}$. This would result in a quadratic dependence of the runtime on the size of the hidden state space, implying a quadratic dependence on the number of haplotypes in the trunk. To this end, define

$$Q[i_{l-1}, \omega_{l-1}] := \sum_{x_{l-1} \in \mathbf{n}_{\omega_{l-1}}} F_{l-1}(i_{l-1}, \omega_{l-1}, x_{l-1}), \tag{52}$$

and

$$R[i_l, \omega_l] := \sum_{\substack{i_{l-1} \in \mathcal{E}, \\ \omega_{l-1} \in \Gamma_{i_{l-1}}}} z(i_l, \omega_l | i_{l-1}, \omega_{l-1}) Q[i_{l-1}, \omega_{l-1}]. \tag{53}$$

Pre-computing $R[i_l, \omega_l]$ and re-using it in (49) allows for an implementation whose runtime only depends linearly on the number of haplotypes in the trunk. Thus, the algorithm to compute the forward probabilities and ultimately the likelihood has runtime complexity $O(Lnd^2)$, where $d = Eg$. Recall that $L$ denotes the number of loci, $n$ the number of haplotypes, $E$ the number of discretization intervals, and $g$ the number of sub-populations at present.

**2.2. Backward algorithm.** The backward probability

$$B_l(s_l) = \mathbb{P}\{H_\Theta^{\alpha,\mathbf{n}}[l+1:L] = h[l+1:L] | T_l^A \in I_{i_l}, G_l = \omega_l, X_l = x_l\} \tag{54}$$

is the probability of observing the alleles $h[l+1:L]$ following locus $l$, conditional on the hidden state at locus $l$. This quantity can again be used to compute the observation probability, but it is also necessary for the expectation-maximization procedure that will be introduced in Section 3. It is possible to write down an explicit backward algorithm for the computation, however, here we proceed along a different route.

To this end define

$$\begin{aligned} F_l^*(s_l) &:= \mathbb{P}\{H_\Theta^{\alpha,\mathbf{n}}[1:(L-l+1)] = h_{[L:l]}, S_{L-l+1} = s_l\} \\ &= \mathbb{P}\{H_\Theta^{\alpha,\mathbf{n}}[l:L] = h_{[L:l]}, S_L = s_l\} \\ &= \mathbb{P}\{H_\Theta^{\alpha,\mathbf{n}}[l:L] = h_{[l:L]}, S_l = s_l\}, \end{aligned} \tag{55}$$

where $h[L:l]$ denotes the reversed vector $(h[L], \ldots, h[l])$, and equality in (55) holds since the transition probability is reversible with respect to the initial distribution. Note that (49) can also be used to compute $F_l^*$, if $F_l$ is replaced by $F_{l-1}^*$, $F_{l-1}$ replaced by $F_l^*$, and the observed alleles are adjusted accordingly to the reversed haplotype $h[L:l]$. Using the modified forward probability $F_l^*$, the backward probability can be obtained via

$$B_l(s_l) = \frac{F_l^*(s_l)}{\xi_\theta(h_l|s_l)\frac{1}{n_{\omega_l}}u(i_l, \omega_l)}. \tag{56}$$

### 3. Parametric Inference via EM

We now present several ways of combining the CSDs introduced in the previous sections in suitable composite likelihood frameworks. We then detail the application of the Expectation Maximization (EM) algorithm to infer demographic parameters in each of these frameworks.

**3.1. Composite Likelihoods.** Assume that the haplotypes in a given sample configuration $\mathbf{n}$ are ordered by enumerating them from 1 to $n$. Thus, $x_i$ denotes the $i$-th haplotype and $\alpha_i$ denotes the sub-population that the $i$-th haplotype resides in at the time the sample is taken, with $1 \leq i \leq n$. Furthermore, for a given permutation $\sigma$ of $\{1, \ldots, n\}$, define

$$\sigma(i, \mathbf{n}) := \sum_{j=1}^{i} \mathbf{e}_{\alpha_{\sigma(j)}, x_{\sigma(j)}} \tag{57}$$

to be the configuration induced by $\sigma$ and a given index $i$, where $\mathbf{e}_{\alpha,x}$ again denotes the configuration with a single haplotype $x$ in sub-population $\alpha$. Further, let

$$\mathbf{n}_{-i} := \mathbf{n} - \mathbf{e}_{\alpha_i,x_i} \quad [58]$$

denote the configuration where haplotype $i$ is removed. As before, denote by $H_\Theta^{\alpha,\mathbf{n}}$ the random additionally sampled haplotype distributed according to the CSD $\pi_\Theta^D(\cdot|\alpha,\mathbf{n})$.

With this notation the product of approximate conditionals (PAC) composite likelihood (7) is given by

$$\mathrm{PAC}_\Theta(\mathbf{n}) := \frac{1}{K}\sum_{\sigma\in\Pi}\prod_{i=1}^{n}\mathbb{P}\{H_\Theta^{\alpha_{\sigma(i)},\sigma(i-1,\mathbf{n})} = x_{\sigma(i)}\}, \quad [59]$$

where $\Pi := \{\sigma_1,\ldots,\sigma_K\}$ are $K$ random permutations of $\{1,\ldots,n\}$. Note that if we would substitute the true CSD in equation (59), each summand would yield the true likelihood of the sample, independent of the ordering $\sigma$. However, if an approximate CSD is used in this formula, the value of the product will depend on the order. This fact had already been noticed by Li and Stephens (7). To mediate the influence of the haplotype-order, following Li and Stephens, we average the product over several random permutations of the ordering.

Replacing the arithmetic mean in definition (59) by a geometric mean yields

$$\mathrm{SuperPAC}_\Theta(\mathbf{n}) := \sqrt[K]{\prod_{\sigma\in\Pi}\prod_{i=1}^{n}\mathbb{P}\{H_\Theta^{\alpha_{\sigma(i)},\sigma(i-1,\mathbf{n})} = x_{\sigma(i)}\}}, \quad [60]$$

another approximation to the sampling probability which we term SuperPAC. Note that the latter also yields the true likelihood if the true CSD would be used instead of an approximation.

The approximate CSD $\pi_\Theta^D(\cdot|\cdot)$ can also be employed in a leave-one-out composite likelihood (LCL)

$$\mathrm{LCL}_\Theta(\mathbf{n}) := \prod_{i=1}^{n}\mathbb{P}\{H_\Theta^{\alpha_i,\mathbf{n}_{-i}} = x_i\}, \quad [61]$$

evaluating the product of all CSDs obtained by leaving each haplotype in turn out of the trunk, or a pairwise composite likelihood (PCL)

$$\mathrm{PCL}_\Theta(\mathbf{n}) := \prod_{i\neq j}\mathbb{P}\{H_\Theta^{\alpha_i,\mathbf{e}_{\alpha_j,x_j}} = x_i\}, \quad [62]$$

consisting of the product of CSDs between all pairs of haplotypes.

**3.2. Objective Functions.** Since the composite likelihoods introduced in the previous paragraphs are combinations of the HMMs underlying the different CSDs, they all comprise of observed random variables $H_\Theta^{\alpha,\mathbf{n}}$, the additionally sampled haplotypes, and latent random variables $S_\Theta^{\alpha,\mathbf{n}}$, the associated sequences of hidden states. To obtain a maximum composite likelihood estimate of the demographic parameters $\Theta$ that best describe a given sample of haplotypes $\mathbf{n}$ under a certain composite likelihood, we apply the standard expectation-maximization (EM) framework (8).

The general outline of the EM algorithm is as follows. Suppose we have parameters $\Theta$, and random variables $\mathbb{X}_\Theta, \mathbb{S}_\Theta$, where $\mathbb{X}_\Theta = \mathbb{X}$ is observed, and $\mathbb{S}_\Theta$ is unobserved (hidden). We would like to find the value of $\Theta$ that maximizes the likelihood $\mathbb{L}(\Theta) = \mathbb{P}(\mathbb{X}_\Theta = \mathbb{X})$. To do so, first choose initial parameters $\Theta^{(0)}$, and then update them iteratively. At step $k+1$, the parameters $\Theta^{(k+1)}$ are obtained by maximizing a certain objective function $Q$ based on $\Theta^{(k)}$, that is

$$\Theta^{(k+1)} = \underset{\Theta}{\mathrm{argmax}}\, Q(\Theta|\Theta^{(k)}). \quad [63]$$

where $Q(\Theta|\Theta^{(k)}) = \mathbb{E}_{\mathbb{S}_{\Theta^{(k)}}}[\log \mathbb{P}(\mathbb{X}_\Theta = \mathbb{X}, \mathbb{S}_\Theta = \mathbb{S}_{\Theta^{(k)}}) \mid \mathbb{X}_{\Theta^{(k)}} = \mathbb{X}]$, where the expectation is taken over $\mathbb{S}_{\Theta^{(k)}}$, as indicated by the subscript. Then $\Theta^{(k)}$ is guaranteed to converge to a local maximum of the likelihood surface $\mathbb{L}(\Theta)$.

We can apply EM to find local maxima of our composite likelihoods $\mathrm{PAC}_\Theta(\mathbf{n})$, $\mathrm{SuperPAC}_\Theta(\mathbf{n})$, $\mathrm{LCL}_\Theta(\mathbf{n})$, $\mathrm{PCL}_\Theta(\mathbf{n})$. To do so, for each composite likelihood, we construct a generative model and random variables $\mathbb{X}_\Theta, \mathbb{S}_\Theta$, such that the composite likelihood is equal to $\mathbb{P}(\mathbb{X}_\Theta = \mathbb{X})$. We then derive $Q_\cdot(\Theta|\Theta^{(k)})$ for each such model.

Note that it is in general not possible to solve the maximization problem in (63) analytically. Thus, in the remainder of this section, we will describe how to evaluate the objective functions for given $\Theta$ and $\Theta^{(k)}$, and employ it in a numerical framework, like the Nelder-Mead simplex algorithm (9), to find a local maximum. The EM framework guarantees that the overall likelihood of the data increases with each parameter update.

**PAC.** Fixing the set of random permutations $\Pi$, definition (59) can be interpreted as a mixture model: First, pick a permutation $\Psi$ uniformly at random from the pool $\Pi$. Then, conditional on $\Psi = \sigma$, generate a random sample $\mathfrak{N}^\sigma_\Theta$: First, sample a haplotype in sub-population $\alpha_{\sigma(1)}$ given an empty trunk. Each allele at each locus is sampled from the stationary distribution of the mutation matrix $P$. Then, sample a second haplotype in sub-population $\alpha_{\sigma(2)}$ given the first haplotype as the already observed trunk; a third haplotype in sub-population $\alpha_{\sigma(3)}$ given the first two in the trunk; and so forth, until a sample of size $n$ is generated. The event that the sample $\mathbf{n}$ is generated in this way is given by

$$\{\mathfrak{N}^\sigma_\Theta = \mathbf{n}\} = \bigcap_{i=1}^{n} \{H_\Theta^{\alpha_{\sigma(i)},\sigma(i-1,\mathbf{n})} = x_{\sigma(i)}\}, \qquad [64]$$

Finally, $\mathrm{PAC}_\Theta(\mathbf{n}) = \mathbb{P}\{\mathfrak{N}^\Psi_\Theta = \mathbf{n}\}$ gives the likelihood of observing the configuration $\mathbf{n}$ under this mixture model, and is equal to (59). Using our previous notation, we have the observed variable $\mathbb{X}_\Theta = \mathfrak{N}^\Psi_\Theta$, and the hidden latent variable $\mathbb{S}_\Theta = \{\Psi, S^{\cdot,\cdot}_\Theta\}$, where $S^{\cdot,\cdot}_\Theta$ is the sequence of hidden states for every CSD.

Let $\Upsilon$ be the random permutation associated with $\Theta^{(k)}$, so $\mathbb{S}_{\Theta^{(k)}} = \{\Upsilon, S^{\cdot,\cdot}_{\Theta^{(k)}}\}$. Then we have

$$\begin{aligned}
&Q_{\mathrm{PAC}}(\Theta|\Theta^{(k)}) \\
&= \mathbb{E}\Big[\log\Big(\mathbb{P}\{\Psi = \Upsilon\} \prod_{i=1}^{n} \mathbb{P}\Big\{S_\Theta^{\alpha_{\Psi(i)},\Psi(i-1,\mathbf{n})} = S_{\Theta^{(k)}}^{\alpha_{\Upsilon(i)},\Upsilon(i-1,\mathbf{n})}, H_\Theta^{\alpha_{\Psi(i)},\Psi(i-1,\mathbf{n})} = x_{\Upsilon(i)} \Big| \Psi = \Upsilon\Big\}\Big) \Big| \mathfrak{N}^\Upsilon_{\Theta^{(k)}} = \mathbf{n}\Big] \\
&= -\log(K) + \sum_{\sigma\in\Pi} \mathbb{P}\{\Upsilon = \sigma | \mathfrak{N}^\Upsilon_{\Theta^{(k)}} = \mathbf{n}\} \\
&\qquad \times \sum_{i=1}^{n} \mathbb{E}\Big[\log \mathbb{P}\Big\{S_\Theta^{\alpha_{\sigma(i)},\sigma(i-1,\mathbf{n})} = S_{\Theta^{(k)}}^{\alpha_{\sigma(i)},\sigma(i-1,\mathbf{n})}, H_\Theta^{\alpha_{\sigma(i)},\sigma(i-1,\mathbf{n})} = x_{\sigma(i)}\Big\} \Big| H_{\Theta^{(k)}}^{\alpha_{\sigma(i)},\sigma(i-1,\mathbf{n})} = x_{\sigma(i)}\Big] \\
&= -\log(K) + \sum_{\sigma\in\Pi} \frac{\mathbb{P}\{\mathfrak{N}^\sigma_{\Theta^{(k)}} = \mathbf{n} | \Upsilon = \sigma\}}{\sum_{\tau\in\Pi} \mathbb{P}\{\mathfrak{N}^\tau_{\Theta^{(k)}} = \mathbf{n} | \Upsilon = \tau\}} \sum_{i=1}^{n} Q_{x_{\sigma(i)}}^{\alpha_{\sigma(i)},\sigma(i-1,\mathbf{n})}(\Theta|\Theta^{(k)}).
\end{aligned} \qquad [65]$$

The second equality follows from partitioning the conditional expectation with respect to $\{\Upsilon = \sigma\}$ and the fact that $\mathbb{P}\{\Psi = \sigma\} = 1/K$. The third equality follows from an application of Bayes' rule and using the definition

$$Q_x^{\alpha,\mathbf{n}}(\Theta|\Theta^{(k)}) := \mathbb{E}\Big[\log \mathbb{P}\{S_\Theta^{\alpha,\mathbf{n}} = S_{\Theta^{(k)}}^{\alpha,\mathbf{n}}, H_\Theta^{\alpha,\mathbf{n}} = x\} \Big| H_{\Theta^{(k)}}^{\alpha,\mathbf{n}} = x\Big]; \qquad [66]$$

the objective function for a single HMM.

**SuperPAC.** Here the generating model is as follows. Again, fix the random set of permutations $\Pi$, but instead of sampling a dataset for a single random permutation as in the PAC mixture model, we obtain $\mathbb{X}_\Theta$ by independently sampling a dataset $\mathfrak{N}^\sigma_\Theta$ for every permutation $\sigma$. We then have $\mathrm{SuperPAC}_\Theta(\mathbf{n})^K = \mathbb{P}(\mathbb{X}_\Theta = (\mathbf{n}, \mathbf{n}, \ldots, \mathbf{n}))$, the likelihood of observing $\{\mathfrak{N}^\sigma_\Theta = \mathbf{n}\}$ for each of the $K$ permutations. The hidden latent variable $\mathbb{S}_\Theta$ is given by the sequence of hidden states for every CSD. The objective function for the

SuperPAC composite likelihood (60) is given by

$$Q_{\mathrm{SuperPAC}}(\Theta|\Theta^{(k)}) = \sum_{\sigma\in\Pi}\sum_{i=1}^{n} Q_{x_{\sigma(i)}}^{\alpha_{\sigma(i)},\sigma(i-1,\mathbf{n})}(\Theta|\Theta^{(k)}), \tag{67}$$

where taking the root can be omitted, since it is a monotone function.

**LCL.** In the LCL (61) case, the objective function is

$$Q_{\mathrm{LCL}}(\Theta|\Theta^{(k)}) = \sum_{i=1}^{n} Q_{x_i}^{\alpha_i,\mathbf{n}_{-i}}(\Theta|\Theta^{(k)}), \tag{68}$$

which is obtained by constructing a generative model where we independently sample the haplotype for each leave-one-out model.

**PCL.** Lastly, the objective function for PCL (62) is

$$Q_{\mathrm{PCL}}(\Theta|\Theta^{(k)}) = \sum_{i\neq j} Q_{x_i}^{\alpha_i,\mathbf{e}_{\alpha_j},x_j}(\Theta|\Theta^{(k)}), \tag{69}$$

which is obtained by a generative model where we independently sample the additional haplotype for each pair.

Equations (65), (67), (68), and (69) show that for each of the composite likelihoods considered here the objective function can be written in terms of the objective functions $Q_{\cdot}^{\cdot,\cdot}$ for the individual HMMs involved. For a general $h$, $\alpha$, and $\mathbf{n}$, this function can be further simplified to obtain

$$\begin{aligned}
Q_h^{\alpha,\mathbf{n}}(\Theta|\Theta^{(k)}) &= \mathbb{E}\Big[\log \mathbb{P}\{S_\Theta^{\alpha,\mathbf{n}} = S_{\Theta^{(k)}}^{\alpha,\mathbf{n}}, H_\Theta^{\alpha,\mathbf{n}} = h\}\Big| H_{\Theta^{(k)}}^{\alpha,\mathbf{n}} = h\Big] \\
&= \sum_{s\in\mathcal{S}} \mathbb{E}\Big[\log\Big((\nu_\Theta(s))^{\mathbb{1}_{\{(S_{\Theta^{(k)}}^{\alpha,\mathbf{n}})_1 = s\}}}\Big)\Big| H_{\Theta^{(k)}}^{\alpha,\mathbf{n}} = h\Big] \\
&\quad + \sum_{s,s'\in\mathcal{S}} \mathbb{E}\Big[\log\Big((\phi_\Theta(s'|s))^{\#\{s\to s'\}}\Big)\Big| H_{\Theta^{(k)}}^{\alpha,\mathbf{n}} = h\Big] \\
&\quad + \sum_{s\in\mathcal{S}}\sum_{a\in E} \mathbb{E}\Big[\log\Big((\xi_\Theta(a|s))^{\#\{s\uparrow a\}}\Big)\Big| H_{\Theta^{(k)}}^{\alpha,\mathbf{n}} = h\Big] \\
&= \sum_{s\in\mathcal{S}} \log(\nu_\Theta(s))\mathbb{P}\{(S_{\Theta^{(k)}}^{\alpha,\mathbf{n}})_1 = s | H_{\Theta^{(k)}}^{\alpha,\mathbf{n}} = h\} \\
&\quad + \sum_{s,s'\in\mathcal{S}} \log(\phi_\Theta(s'|s))\mathbb{E}[\#\{s\to s'\}|H_{\Theta^{(k)}}^{\alpha,\mathbf{n}} = h] \\
&\quad + \sum_{i\in\mathcal{E},\omega\in\Gamma_i}\sum_{a,t\in E} \log(\xi_\Theta(a|i,\omega,t))\mathbb{E}[\#\{(i,\omega,t)\uparrow a\}|H_{\Theta^{(k)}}^{\alpha,\mathbf{n}} = h].
\end{aligned} \tag{70}$$

The initial $\nu$, transition $\phi$, and emission $\xi$ probabilities are given in (28), (41), and (47). Here the subscripts $\Theta$ is used to emphasize their dependence on the demographic parameters. Furthermore, $\#\{s\uparrow a\}$ denotes the number of times allele $a$ is emitted from hidden state $s$ for a given realization of $S_{\Theta^{(k)}}^{\alpha,\mathbf{n}}$ and $H_{\Theta^{(k)}}^{\alpha,\mathbf{n}}$, and $\#\{s\to s'\}$ is the number of transitions from hidden state $s$ to $s'$. Note that we slightly abuse the notation by conditioning on the trunk allele instead of the trunk haplotype in the emission probability $\xi$ on the last line. We adjust the number of emissions appropriately.

The second summand on the right hand side of (70) (the transition part) can be further modified to

$$\begin{aligned}
&\sum_{i\in\mathcal{E},\omega\in\Gamma_i}\sum_{i'\in\mathcal{E},\omega'\in\Gamma_{i'}}\sum_{x\in\mathbf{n}_\omega,x'\in\mathbf{n}_{\omega'}}\log\big(y_\Theta(i,\omega)\delta_{i,i'}\delta_{\omega,\omega'}\delta_{x,x'}+\frac{1}{n_{\omega'}}z_\Theta(i',\omega'|i,\omega)\big)\\
&\qquad\qquad\times\mathbb{E}\Big[\#\{(i,\omega,x)\to(i',\omega',x')\}\Big|H^{\alpha,\mathbf{n}}_{\Theta^{(k)}}=h\Big]\\
&\quad=\sum_{i\in\mathcal{E},\omega\in\Gamma_i}\sum_{i'\in\mathcal{E},\omega'\in\Gamma_{i'}}\log\big(\frac{1}{n_{\omega'}}z_\Theta(i',\omega'|i,\omega)\big)\Big(\mathbb{E}\Big[\#\{(i,\omega)\to(i',\omega')\}\Big|H^{\alpha,\mathbf{n}}_{\Theta^{(k)}}=h\Big]\\
&\qquad\qquad-\delta_{i,i'}\delta_{\omega,\omega'}\sum_{x\in\mathbf{n}_\omega}\mathbb{E}\Big[\#\{(i,\omega,x)\to(i,\omega,x)\}\Big|H^{\alpha,\mathbf{n}}_{\Theta^{(k)}}=h\Big]\Big)\\
&\qquad+\sum_{i\in\mathcal{E},\omega\in\Gamma_i}\log\big(y_\Theta(i,\omega)+\frac{1}{n_\omega}z_\Theta(i,\omega|i,\omega)\big)\sum_{x\in\mathbf{n}_\omega}\mathbb{E}\Big[\#\{(i,\omega,x)\to(i,\omega,x)\}\Big|H^{\alpha,\mathbf{n}}_{\Theta^{(k)}}=h\Big],
\end{aligned}\tag{71}$$

with

$$\#\{(i,\omega)\to(i',\omega')\}:=\sum_{x\in\mathbf{n}_\omega,x'\in\mathbf{n}_{\omega'}}\#\{(i,\omega,x)\to(i',\omega',x')\}.\tag{72}$$

We introduce this modification, since a naïve implementation of the left hand side of (71) would depend quadratically on the number of haplotypes in the trunk, whereas the right hand side only depends linearly on this number.

**3.3. Computing the Conditional Expectations.** We now provide the details on how to compute the conditional probabilities and expectations that are required to evaluate (71) and the objective function (70), which can then be used to evaluate the objective functions for the different composite likelihoods. Assume that for all $l\in\{1,\dots,L\}$ and all $s\in\mathcal{S}$ the forward probabilities $F_l(s)$ and the backward probabilities $B_l(s)$ introduced in Section 2 haven been computed under the parameters $\Theta^{(k)}$.

The posterior probabilities for the initial hidden state are then given by

$$\begin{aligned}
\mathbb{P}\{(S^{\alpha,\mathbf{n}}_{\Theta^{(k)}})_1=s|H^{\alpha,\mathbf{n}}_{\Theta^{(k)}}=h\}&=\frac{\mathbb{P}\{(S^{\alpha,\mathbf{n}}_{\Theta^{(k)}})_1=s,H^{\alpha,\mathbf{n}}_{\Theta^{(k)}}=h\}}{\mathbb{P}\{H^{\alpha,\mathbf{n}}_{\Theta^{(k)}}=h\}}\\
&=\frac{1}{\mathbb{P}\{H^{\alpha,\mathbf{n}}_{\Theta^{(k)}}=h\}}\sum_{s\in\mathcal{S}}\nu_{\Theta^{(k)}}(s)\xi_{\Theta^{(k)}}(h_1|s)B_1(s).
\end{aligned}\tag{73}$$

The conditional expectation in (71) that is marginalized over the absorbing haplotypes can be evaluated using

$$\begin{aligned}
&\mathbb{E}\Big[\#\{(i,\omega)\to(i',\omega')\}\Big|H^{\alpha,\mathbf{n}}_{\Theta^{(k)}}=h\Big]\\
&=\frac{1}{\mathbb{P}\{H^{\alpha,\mathbf{n}}_{\Theta^{(k)}}=h\}}\sum_{l=1}^{L}\sum_{x\in\mathbf{n}_\omega,x'\in\mathbf{n}_{\omega'}}\big(y_{\Theta^{(k)}}(i,\omega)\delta_{i,i'}\delta_{\omega,\omega'}\delta_{x,x'}+\frac{1}{n_{\omega'}}z_{\Theta^{(k)}}(i',\omega'|i,\omega)\big)\\
&\qquad\qquad\times F_l(i,\omega,x)\xi_{\Theta^{(k)}}(h_{l+1}|i',\omega',x')B_{l+1}(i',\omega',x')\\
&=\frac{1}{\mathbb{P}\{H^{\alpha,\mathbf{n}}_{\Theta^{(k)}}=h\}}\sum_{l=1}^{L}\Big[\frac{1}{n_{\omega'}}z_{\Theta^{(k)}}(i',\omega'|i,\omega)\Big(\sum_{x\in\mathbf{n}_\omega}F_l(i,\omega,x)\Big)\times\sum_{x'\in\mathbf{n}_{\omega'}}\xi_{\Theta^{(k)}}(h_{l+1}|i',\omega',x')B_{l+1}(i',\omega',x')\\
&\qquad\qquad+\delta_{i,i'}\delta_{\omega,\omega'}y_{\Theta^{(k)}}(i,\omega)\sum_{x\in\mathbf{n}_\omega}F_l(i,\omega,x)\xi_{\Theta^{(k)}}(h_{l+1}|i,\omega,x)B_{l+1}(i,\omega,x)\Big].
\end{aligned}\tag{74}$$

Again, the computation of right hand side only depends linearly on the number of haplotypes in the trunk.

The expectation involving the transition from a certain hidden state to itself is given by

$$\mathbb{E}\Big[\#\{(i,\omega,x)\to(i,\omega,x)\}\Big|H^{\alpha,\mathbf{n}}_{\Theta^{(k)}}=h\Big] \\ =\frac{1}{\mathbb{P}\{H^{\alpha,\mathbf{n}}_{\Theta^{(k)}}=h\}}\big(y_{\Theta^{(k)}}(i,\omega)+\frac{1}{n_{\omega'}}z_{\Theta^{(k)}}(i',\omega'|i,\omega)\big)\sum_{l=1}^{L}F_l(i,\omega,x)\xi_{\Theta^{(k)}}(h_{l+1}|i,\omega,x)B_{l+1}(i,\omega,x). \quad [75]$$

Finally,

$$\mathbb{E}[\#\{(i,\omega,t)\uparrow a\}|H^{\alpha,\mathbf{n}}_{\Theta^{(k)}}=h]=\frac{1}{\mathbb{P}\{H^{\alpha,\mathbf{n}}_{\Theta^{(k)}}=h\}}\sum_{l=1}^{L}\mathbb{1}_{\{h_l=a\}}\sum_{\substack{x\in\mathbf{n}_\omega\\ x_l=t}}F_l(i,\omega,x)B_l(i,\omega,x) \quad [76]$$

gives the conditional expectation of the number of emissions of a certain type. The time complexity to evaluate the objective function (70) is $O(nd^2)$, where $d=Eg$. The overall complexity for the EM algorithm depends on the particular composite likelihood that is used.

### 4. Improving computational efficiency

We now introduce two modifications in order to speed up the computations of the forward-backward algorithm. The runtimes will still depend linearly on the number of loci, but the number of loci that effectively have to be considered will be reduced. In what follows, assume that the demographic parameters $\Theta$, an additional haplotype $h$, a corresponding additional sub-population $\alpha$, and a configuration of trunk haplotypes $\mathbf{n}$ are given, and consider the computations for the CSD $\pi^D_\Theta(h|\alpha,\mathbf{n})=\mathbb{P}\{H^{\alpha,\mathbf{n}}_\Theta=h\}$.

**4.1. Locus skipping.** First we will detail a modification that decreases the number of effective loci by "skipping" over non-polymorphic loci. A similar modification has been introduced before (10) and it requires that the mutation matrix is such that every allele mutates to every other allele at the same rate. For example, this requirement is satisfied by the mutation matrix with $P_{a,a'}=\frac{1}{|E|-1}$ if $a\neq a'$, and $P_{a,a}=0$. It follows that

$$\xi(a|i,\omega,a)=\xi(a'|i,\omega,a') \quad [77]$$

holds for all $a,a'\in E$, $i\in\mathcal{E}$, and $\omega\in\Gamma_i$. The modified computations produce the exact result if the given mutation matrix satisfies this requirement. However, even if this is not the case, the computational benefit might outweigh the approximation error. Furthermore, define the set of non-polymorphic loci by

$$\mathscr{N}:=\{1\leq l\leq L|h[l]=x[l],\forall x\in\mathbf{n}\}, \quad [78]$$

that is, the set of all loci, where the additional haplotype and all the trunk haplotypes carry the same allele.

Then, given two hidden states $s=(i,\omega,x),s'=(i',\omega',x')\in\mathcal{S}$, define the $k$-step transition probability as

$$\phi^{(k)}(s'|s) \\ :=\mathbb{P}\{T^A_{l+k}\in I_{i'},G_{l+k}=\omega',X_{l+k}=x'|T^A_l\in I_i,G_l=\omega,X_l=x,\{l+1,\dots,l+k-1\}\subset\mathscr{N}\}. \quad [79]$$

By conditioning on $\{l+1,\dots,l+k-1\}\subset\mathscr{N}$ we include the requirement that the intervening loci $\{l+1,\dots,l+k-1\}$ are not polymorphic. The base case is given by

$$\phi^{(1)}(s'|s)=y^{(1)}(i,\omega)\delta_{i',i}\delta_{\omega',\omega}\delta_{x',x}+z^{(1)}(i',\omega'|i,\omega)\frac{1}{n_{\omega'}} \\ =y(i,\omega)\delta_{i',i}\delta_{\omega',\omega}\delta_{x',x}+z(i',\omega'|i,\omega)\frac{1}{n_{\omega'}}=\phi(s'|s), \quad [80]$$

the transition probability defined in (41). Now for a given $k$, and any $k_1, k_2 \in \mathbb{N}$ with $k_1 + k_2 = k$, the recursive relation

$$\begin{aligned}\phi^{(k)}(s'|s) &= \sum_{t \in \mathcal{S}} \phi^{(k_2)}(s'|t)\xi(h_{l+k_1}|t)\phi^{(k_1)}(t|s) \\ &= \sum_{j \in \mathcal{E}, \psi \in \Gamma_j} \xi(a|j,\psi,a) \sum_{v \in \mathbf{n}_\psi} \phi^{(k_2)}(s'|j,\psi,v)\phi^{(k_1)}(j,\psi,v|s) \\ &= y^{(k)}(i,\omega)\delta_{i',i}\delta_{\omega',\omega}\delta_{x',x} + z^{(k)}(i',\omega'|i,\omega)\frac{1}{n_{\omega'}}\end{aligned} \quad [81]$$

holds, with

$$y^{(k)}(i,\omega) := \xi(a|i,\omega,a)y^{(k_2)}(i,\omega)y^{(k_1)}(i,\omega), \quad [82]$$

and

$$\begin{aligned}z^{(k)}(i',\omega'|i,\omega) := {}& \xi(a|i,\omega,a)z^{(k_2)}(i',\omega'|i,\omega)y^{(k_1)}(i,\omega) \\ &+ \xi(a|i',\omega',a)y^{(k_2)}(i',\omega')z^{(k_1)}(i',\omega'|i,\omega) \\ &+ \sum_{j \in \mathcal{E}, \psi \in \Gamma_j} \xi(a|j,\psi,a)z^{(k_2)}(i',\omega'|j,\psi)z^{(k_1)}(j,\psi|i,\omega).\end{aligned} \quad [83]$$

The $k$-step transition probabilities can be employed as follows. Denote by $\mathscr{L}' := \{1\} \cup \overline{\mathscr{N}} \cup \{L\}$ the set of polymorphic loci plus the first and the last. Further, define

$$\mathfrak{p}(l,l') := \{\mathfrak{n}(l,l')\} \cup \mathfrak{p}(\mathfrak{n}(l,l'),l'), \quad [84]$$

with $\mathfrak{n}(l,l') := \max\{l + 2^m | m \in \mathbb{N}_0, l + 2^m < l'\}$ for $l + 1 < l'$, and $\mathfrak{p}(l, l+1) := \emptyset$. Now

$$\mathscr{L} := \mathscr{L}' \cup \bigcup_{(l,l') \text{ consecutive in } \mathscr{L}'} \mathfrak{p}(l,l') \quad [85]$$

is the set of polymorphic loci, plus a scaffold that guarantees that the distance between two consecutive loci in $\mathscr{L}$ is always a power of 2. Further, every locus between 1 and $L$ that is not an element of $\mathscr{L}$ is guaranteed to be non-polymorphic. Thus, the forward $F_l(s)$ and backward $B_l(s)$ probabilities can be computed for $l \in \mathscr{L}$ using only transition probabilities of the form $\phi^{(k)}$ given in (79) with $k = 2^m$, where $m$ is an non-negative integer from 0 to the maximal exponent needed for the possible steps in $\mathscr{L}$. The initial and emission probabilities do not need to be modified.

Previously, since all steps along the sequence in the EM algorithm detailed in Section 3 have the same size, only one term involving the transition probability occurs on the right hand side of (70). To implement the possibility of different step sizes, steps of the same size have to be grouped together, and a term like the former has to be added for each group. If the set of polymorphic loci $\mathscr{L}'$ would be used directly for the computations, there would in general be a large number of different sizes, and the EM algorithm would not be very efficient. However, by using the set $\mathscr{L}$ instead, it is guaranteed that the sizes of the possible steps are all powers of two, and thus the EM algorithm can still be implemented efficiently.

**4.2. Multi-locus HMM-step handler.** A different approach to reduce the effective number of loci is by combining neighboring loci within a window into "meta"-loci. To this end, assume that a window-size $b \in \mathbb{N}$ is given, and define the set of "meta"-loci as $\mathscr{L}^\star := \{0, \ldots, \lfloor (L-1)/b \rfloor\}$. Mathematically, this approach is equivalent to restricting the hidden states at all loci in the set $\{(l^\star \cdot b) + 1, \ldots, (l^\star \cdot b) + b\}$ to be identical, and setting the recombination rates between $(l^\star \cdot b) + b$ and $((l^\star + 1) \cdot b) + 1$ to $b \cdot \rho$, for each $l^\star \in \mathscr{L}^\star$.

Combining the loci can be implemented as follows. Define modified forward probabilities $F^\star_{l^\star}(s)$ for all $l^\star \in \mathscr{L}^\star$, and compute them according to (49), with modified transition $\phi^\star$ and emission $\xi^\star$ probabilities.

The transition probabilities $\phi^\star$ are essentially given by definition (41), only a recombination rate of $b \cdot \rho$ has to be used instead of $\rho$. At a given locus $l^\star \in \mathscr{L}^\star$, for a given hidden state $s = (i, \omega, x)$, the "allele" of the additional haplotype is $h[(l^\star \cdot b)+1 : (l^\star \cdot b)+b]$ and the "allele" of the trunk haplotype is $x[(l^\star \cdot b)+1 : (l^\star \cdot b)+b]$. Thus, the emission probability at locus $l^\star$ is given by

$$\begin{aligned} &\xi^\star\big(h[(l^\star \cdot b) + 1 : (l^\star \cdot b) + b]\big|i, \omega, x[(l^\star \cdot b) + 1 : (l^\star \cdot b) + b]\big) \\ &\quad = \prod_{l \in \{(l^\star \cdot b)+1, \ldots, (l^\star \cdot b)+b\}} \xi\big(h[l]\big|i, \omega, x[l]\big). \end{aligned} \qquad [86]$$

The initial probabilities $\nu$ remain unchanged. The modified backward probabilities $B^\star_{l^\star}(s)$ and the EM algorithm can be adjusted accordingly.

### 5. Migrating trunk

The unchanging trunk previously described has some drawbacks (also mentioned in the main text). In particular, under the true coalescent, a trunk lineage may absorb with the additional lineage in a sub-population different from the one it resides in at present, due to migration of the trunk lineage. Furthermore, going backwards in time, the rate of absorption of the additional lineage decreases, due to coalescence events within the trunk.

To mitigate these drawbacks, we modify the approximate CSD. Under the model outlined in Section 1, a hidden state $s = (i, \omega, x) \in \mathcal{S}$ represents the event that the additional lineage is absorbed during interval $I_i$, in sub-population $\omega \in \Gamma_i$, into the trunk-lineage $x$. In the modified model, a hidden state $s^\dagger = (i, \omega^\dagger, x) \in \mathcal{E} \times \Gamma \times \mathbf{n}_{\omega^\dagger}$ represents the event that during the interval $I_i$ the lineage of the additional haplotype absorbs into the lineage of the haplotype $x$ that resides in $\omega^\dagger$ at the present.

Now, approximate the genealogy relating the haplotypes in the trunk under the true model as follows. First, recall that the number of absorbing lineages in the trunk determines the absorption rates in definition (4), and they were assumed constant for each sub-population in Section 1. Under the coalescent with migration, these numbers are given by a stochastic process, the ancestral process, that evolves due to coalescence and migration events. Using the full stochastic process is prohibitive, however, Jewett and Rosenberg (11) showed that often times its expected value can be used instead without much loss in accuracy. To this end consider a given epoch $\epsilon \in \mathcal{E}$, with $I_\epsilon = [t_{\epsilon-1}, t_\epsilon)$. The expected number of trunk lineages in each sub-population $\{n_\gamma^{(\epsilon)}(t_{\epsilon-1})\}_{\gamma \in \Gamma_\epsilon}$ at the beginning of epoch $\epsilon$ are given by $\{n_\gamma\}_{\gamma \in \Gamma}$, if $\epsilon = 1$, and

$$\Big\{ \sum_{\substack{\delta \in \Gamma_{\epsilon-1} \\ \delta \subset \gamma}} n_\delta^{(\epsilon-1)}(t_{\epsilon-1}) \Big\}_{\gamma \in \Gamma_\epsilon} \qquad [87]$$

otherwise. The dynamics of the expected number of lineages during epoch $\epsilon$ can be approximated by the system of differential equations

$$\frac{d}{dt} n_\gamma^{(\epsilon)}(t) = -\frac{1}{\kappa_\gamma^{(\epsilon)}} \binom{n_\gamma^{(\epsilon)}(t)}{2} \mathbb{1}_{\{n_\gamma^{(\epsilon)}(t) > 1\}} + \sum_{\substack{\delta=1 \\ \delta \neq \gamma}}^{|\Gamma_\epsilon|} \big(m_{\delta,\gamma}^{(\epsilon)} n_\delta^{(\epsilon)}(t) - m_{\gamma,\delta}^{(\epsilon)} n_\gamma^{(\epsilon)}(t)\big), \qquad [88]$$

for $\gamma \in \Gamma_\epsilon$, c.f. (11, Equation 26). The additional indicator function in the first summand balances out the fact that a term involving the variance is missing in this approximation. For each epoch $\epsilon \in \mathcal{E}$, these differential equations can be solved numerically. Then, replace $n_\gamma$ by $\frac{1}{2}\big(n_\gamma^{(\epsilon)}(t_{\epsilon-1}) + n_\gamma^{(\epsilon)}(t_\epsilon)\big)$ for each $\gamma \in \Gamma_\epsilon$ in (4), and compute $\tilde{u}$, $\tilde{y}$, $\tilde{z}$, and $\tilde{\xi}$ using these modified absorption matrices in (29), (42), (43), and (47), respectively.

To approximate the effect of the migration dynamics in the trunk on the absorption dynamics of the lineage of the additional haplotype, define $p^{(\epsilon)}(\gamma, \delta)$ as the probability that a lineage residing in sub-population $\gamma$ at present resides in sub-population $\delta$ at the beginning of epoch $\epsilon$. Here $\gamma \in \Gamma$ and $\delta \in \Gamma_\epsilon$. Further, let

$q^{(\epsilon)}(\gamma,\delta)$ be the corresponding probability at the end of the epoch. If $\epsilon = 1$, then $p^{(\epsilon)}(\gamma,\delta) = \delta_{\gamma,\delta}$. If $\epsilon > 1$, then

$$p^{(\epsilon)}(\gamma,\delta) = \sum_{\substack{\zeta\in\Gamma_{\epsilon-1}\\ \zeta\subset\delta}} q^{(\epsilon-1)}(\gamma,\zeta) \quad [89]$$

holds. Furthermore,

$$q^{(\epsilon)}(\gamma,\delta) = \sum_{\mu\in\Gamma_\epsilon} p^{(\epsilon)}(\gamma,\mu) \begin{cases} \left(e^{(t_\epsilon - t_{\epsilon-1})M_\epsilon}\right)_{\mu,\delta}, & \text{if } I_\epsilon \neq \emptyset, \\ \left(Y_\epsilon\right)_{\mu,\delta}, & \text{if } I_\epsilon = \emptyset. \end{cases} \quad [90]$$

Lastly, define the average of $p^{(\epsilon)}$ and $q^{(\epsilon)}$, weighted by the number of haplotypes in a certain sub-population, as

$$r^{(\epsilon)}(\gamma,\delta) := \frac{1}{2}\big(p^{(\epsilon)}(\gamma,\delta) + q^{(\epsilon)}(\gamma,\delta)\big)\cdot n_\gamma, \quad [91]$$

and define

$$\tilde{r}^{(\epsilon)}(\gamma,\delta) = \frac{1}{\sum_{\mu\in\Gamma} r^{(\epsilon)}(\mu,\delta)} r^{(\epsilon)}(\gamma,\delta) \quad [92]$$

as the fraction of lineages residing sub-population $\delta$ during epoch $\epsilon$ that reside in sub-population $\gamma$ at present.

Combining the quantities introduced in the previous paragraphs, define the modified initial probabilities as

$$\nu^\dagger(i,\omega^\dagger,x) := \frac{1}{n_{\omega^\dagger}} \underbrace{\sum_{\gamma\in\Gamma_i} \tilde{r}^{(i)}(\omega^\dagger,\gamma)\tilde{u}(i,\gamma)}_{=:\tilde{v}(i,\omega^\dagger)}, \quad [93]$$

that is, the probability of being absorbed during epoch $i$ into lineage $x$ residing in sub-population $\omega^\dagger$ at present is given by considering the probability of being absorbed in a certain sub-population times the fraction of lineages in that sub-population that are ancestral to lineages in sub-population $\omega^\dagger$ at present, and then summing this over all sub-populations. Along similar lines, define the modified transition probabilities

$$\begin{aligned} \phi^\dagger(i',\psi^\dagger,&x'|i,\omega^\dagger,x) \\ &= \frac{1}{\tilde{v}(i,\omega^\dagger)}\Big(\delta_{i',i}\delta_{\psi^\dagger,\omega^\dagger}\delta_{x',x}\sum_{\gamma\in\Gamma_i}\tilde{r}^{(i)}(\omega^\dagger,\gamma)\tilde{y}(i,\gamma)\tilde{u}(i,\gamma) \\ &\qquad + \frac{1}{n_{\psi^\dagger}}\sum_{\delta\in\Gamma_{i'}}\sum_{\gamma\in\Gamma_i}\tilde{r}^{(i')}(\psi^\dagger,\delta)\tilde{r}^{(i)}(\omega^\dagger,\gamma)\tilde{z}(i',\delta|i,\gamma)\tilde{u}(i,\gamma)\Big). \end{aligned} \quad [94]$$

and the modified emission probabilities

$$\xi^\dagger(a|i,\omega^\dagger,t) = \frac{1}{\tilde{v}(i,\omega^\dagger)}\sum_{\gamma\in\Gamma_i}\tilde{r}^{(i)}(\omega^\dagger,\gamma)\tilde{\xi}(a|i,\gamma,t)\tilde{u}(i,\gamma). \quad [95]$$

These probabilities can then be used in appropriately modified versions of the forward, backward, and EM algorithm.

### 6. Decoupling HMM discretization from demographic history

We now describe how to employ a discretization for the HMM computations that differs from the partition induced by the demographic history. To this end, define a partition of the real line into intervals $\{J_j\} := \{[t_{j-1}, t_j)\}$ that is to be used to discretize the HMM. Note that, for convenience, we abuse the subscript notation for $t$ slightly. The hidden states of our HMM are then $(j, \omega^\dagger, x)$, where $j$ is an index of the discretization intervals $\{J_j\}$, and $\omega^\dagger$ and $x$ are as before. We define a third partition

$$\{K_k\} := \bigcup_{\{I_\epsilon\}} \bigcup_{\{J_j\}} \{I_\epsilon \cap J_j\}. \qquad [96]$$

Note that $\{K_k\}$ is a refinement of $\{I_\epsilon\}$ and $\{J_j\}$, that is for all $k$ the inclusion $K_k \subset I_\epsilon$ holds for some $\epsilon$, and $K_k \subset J_j$ for some $j$. In particular, note that the population sizes and migration rates are constant within each refined interval $K_k$. Thus we can work with the "refined" demographic history with epochs $\{K_k\}$ instead of $\{I_\epsilon\}$. Specifically, associate with interval $K_k$ the set of sub-populations $\Gamma_k := \Gamma_\epsilon$ and a migration matrix $M_k := M_\epsilon$, with $\epsilon$ such that $K_k \subset I_\epsilon$. Assign $Y_k$ to the intervals of length zero accordingly.

As in Section 5, compute $\tilde{u}$, $\tilde{y}$, $\tilde{z}$, and $\tilde{\xi}$ according to (29), (42), (43), and (47), respectively, using the modified absorption rates and the discretization $\{K_k\}$. Also, define $\tilde{r}^{(k)}(\gamma, \delta)$ accordingly, using this discretization in (92). Since $\{K_k\}$ is a refinement of $\{J_j\}$, we can then use these quantities to compute the initial $\nu^\dagger$, transition $\phi^\dagger$, and emission $\xi^\dagger$ probabilities for the discretization $\{J_j\}$, analogously to (93), (94), and (95). Specifically, replace $\tilde{r}^{(j)}(\omega^\dagger, \gamma)\tilde{u}(j, \gamma)$ in (93) by

$$\sum_{k:K_k \subset J_j} \tilde{r}^{(k)}(\omega^\dagger, \gamma)\tilde{u}(k, \gamma). \qquad [97]$$

In (95), replace $\tilde{r}^{(j)}(\omega^\dagger, \gamma)\tilde{y}(j, \gamma)\tilde{u}(j, \gamma)$ with

$$\sum_{k:K_k \subset J_j} \tilde{r}^{(k)}(\omega^\dagger, \gamma)\tilde{y}(k, \gamma)\tilde{u}(k, \gamma), \qquad [98]$$

and $\tilde{r}^{(i')}(\psi^\dagger, \delta)\tilde{r}^{(i)}(\omega^\dagger, \gamma)\tilde{z}(i', \delta|i, \gamma)\tilde{u}(i, \gamma)$ with

$$\sum_{k':K_{k'} \subset J_{j'}} \sum_{k:K_k \subset J_j} \tilde{r}^{(k')}(\psi^\dagger, \delta)\tilde{r}^{(k)}(\omega^\dagger, \gamma)\tilde{z}(k', \delta|k, \gamma)\tilde{u}(k, \gamma). \qquad [99]$$

Lastly, replace $\tilde{r}^{(j)}(\omega^\dagger, \gamma)\tilde{\xi}(a|j, \gamma, t)\tilde{u}(j, \gamma)$ in (95) by

$$\sum_{k:K_k \subset J_j} \tilde{r}^{(k)}(\omega^\dagger, \gamma)\tilde{\xi}(a|k, \gamma, t)\tilde{u}(k, \gamma). \qquad [100]$$

Using these probabilities all computations for the HMM, and the algorithms based on them, can be computed using the discretization $\{J_j\}$ independent of the partition $\{I_\epsilon\}$ that is used for the demographic history.

### Supplementary Figures

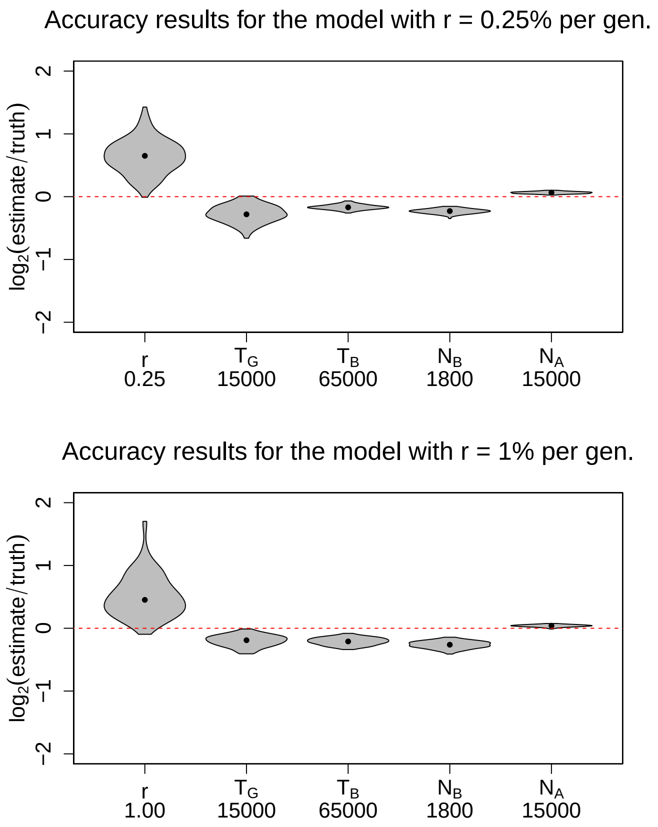


**Fig. S1. Accuracy results of our method, `diCal2`, for the recent exponential growth model shown in Figure 1A of the main text with expansion rate $r = 0.25\%$ and $1.0\%$ per generation.** Parameter estimates were obtained using only 10 haplotypes, which is much less than the sample size (thousands to tens of thousands) required by SFS-based methods to get good estimates. Each violin plot shows the base-2 logarithm of the relative error (estimate/truth) for the analysis of 100 simulated datasets. Thus, a value of 0 corresponds to an exact estimate, whereas $+1$ is a two-fold over- and $-1$ is a two-fold underestimate. True parameter values are shown on the $x$-axis.

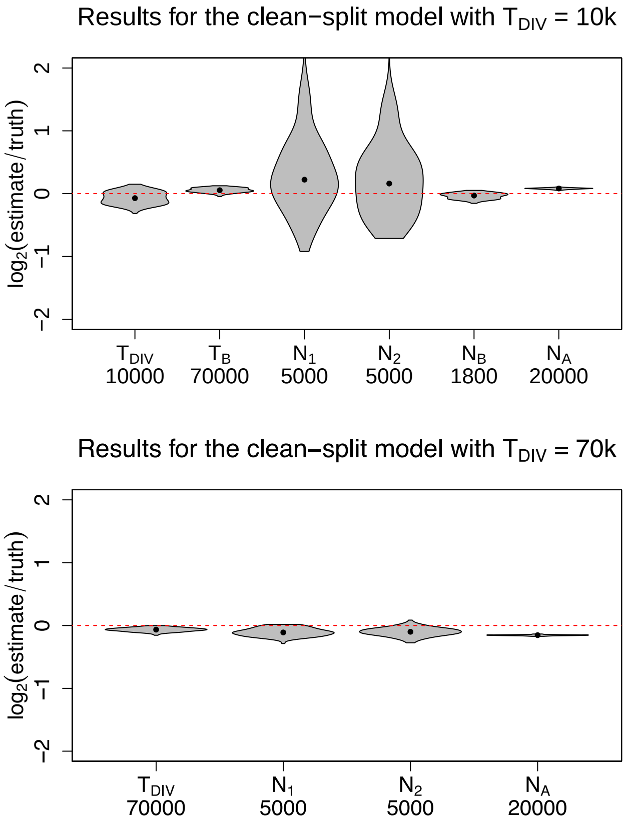


**Fig. S2. Accuracy results for the clean-split model (no gene flow, $m = 0$) shown in Figure 1B of the main text with divergence time $T_{\text{DIV}} = 10$ and $70$ ka.** Using only two haplotypes in each extant population, the parameters of this clean-split model could be estimated very accurately.

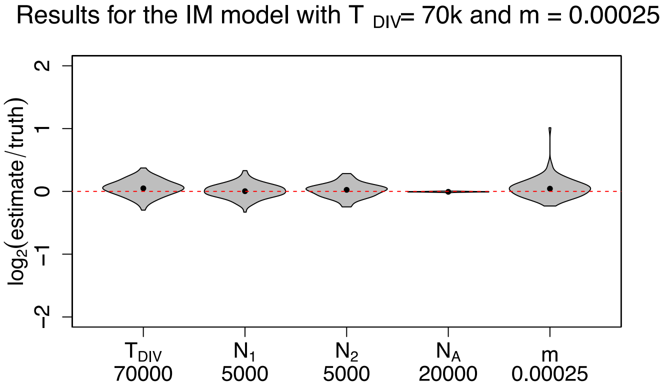


**Fig. S3. Accuracy results for the isolation with migration (IM) model shown in Figure 1B of the main text with divergence time $T_{\text{DIV}} = 70$ ka, and migration probability $m = 0.00025$.** As in the clean-split case, only two haplotypes in each extant population were used. Most parameter estimates show little bias or variability. See the text for further discussion.

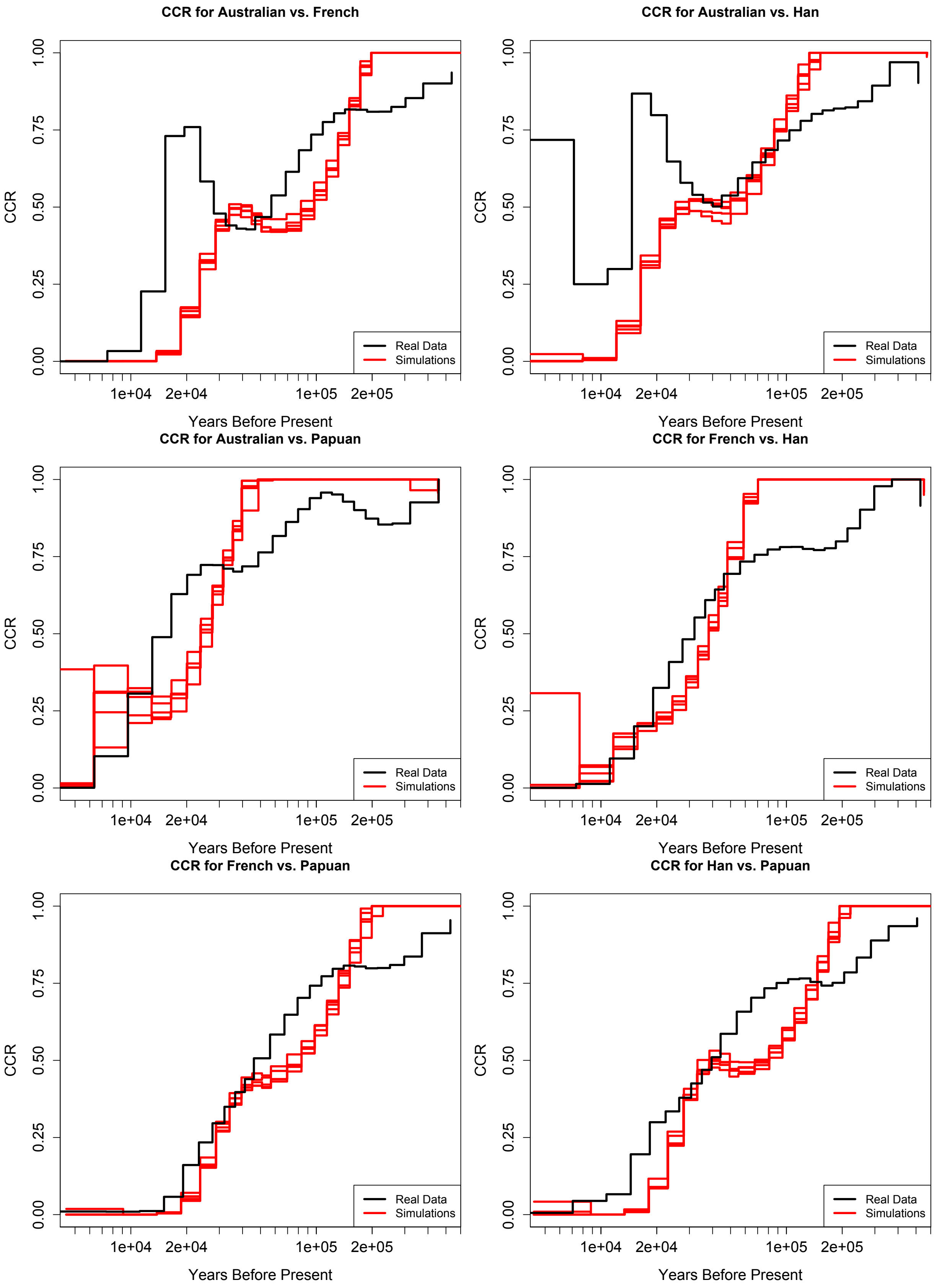


**Fig. S4. Goodness-of-fit evaluated using cross-coalescence rate curves.** For each population pair, we estimated the Cross-coalescence rate curve from the SGDP data (shown in black) and for datasets simulated under the estimated parameters (shown in red).

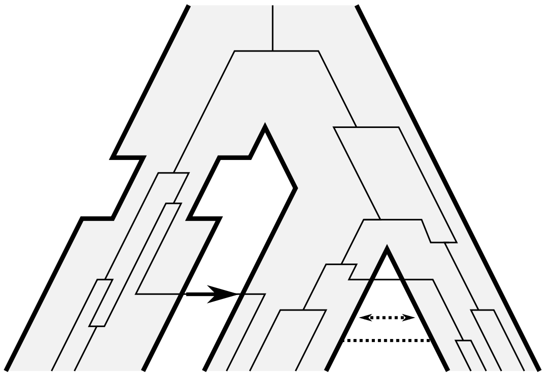

**Fig. S5. An example of a demographic history and a realization of the coalescent with recombination under this history.** In this example, there are $E = 8$ epochs and epoch $3$ has length zero, so $t_2 = t_3$. The demographic structure is given by $\Gamma_1 = \Gamma_2 = \Gamma_3 = \Gamma_4 = \{\{1\}, \{2\}, \{3\}\}$, $\Gamma_5 = \Gamma_6 = \Gamma_7 = \{\{1\}, \{2, 3\}\}$, and $\Gamma_8 = \{\{1, 2, 3\}\}$. All migration rates associated with all epochs but $3$ are zero, except $m^{(2)}_{2,3} = m^{(2)}_{3,2} = m^{(4)}_{2,3} = m^{(4)}_{3,2} = m$ (and the rates on the diagonal accordingly). The instantaneous migration probabilities associated with epoch $3$ are all zero, but $y^{(3)}_{2,1} \geq 0$. The bottleneck is implemented by setting $\kappa^{(6)}_{\gamma_{6,1}} < \kappa^{(5)}_{\gamma_{5,1}} = \kappa^{(7)}_{\gamma_{7,1}}$.

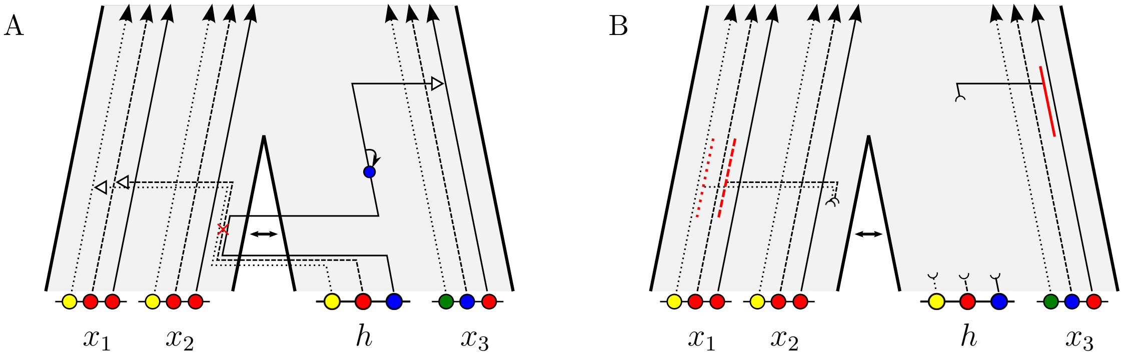


**Fig. S6. Two example realizations of the approximations to the true CSD.** In these examples, the demography $\Theta$ describes an ancestral population that splits into two, with subsequent gene flow. The already observed configuration consists of haplotypes $x_1$ and $x_2$ in the first population and $x_3$ in the second. The additional haplotype $h$ is sampled in the second population. (A) $\pi_\Theta^T\left(h\middle|2,\{x_1,x_2,x_3\}\right)$. The CSD $\pi_\cdot^T(\cdot|\cdot)$ approximates the true genealogy relating the observed haplotypes by an unchanging trunk. The dotted, dashed, and solid lines represent the lineages at locus 1, 2, and 3, respectively. At the first locus, the marginal additional lineage undergoes a migration event and is absorbed into the trunk-lineage of $x_1$. A recombination event, indicated by the red cross, separates the lineages at locus 2 and 3. Thus, up to the time of the breakpoint, the additional lineages are the same. At locus 3, it then undergoes migration independently and is absorbed at a different time into the trunk-lineage of $x_3$. The alleles at each locus are then propagated to the present accounting for possible mutations, indicated by the black arrow. (B) $\pi_\Theta^D\left(h\middle|2,\{x_1,x_2,x_3\}\right)$. Under the approximation $\pi_\cdot^D(\cdot|\cdot)$, the absorbing trunk-lineages at each locus are as before, however, only the intervals (indicated in red) that the absorption times falls into are recorded.

### Supplementary Table

**Table S1. Bootstrap results.** The following table shows the parameter estimates and bootstrap results from the analysis of different pairs of population from the SGDP data using the demographic model in Figure 3 of the main text. For each pair, the table provides the raw estimates, 10 parametric bootstrap estimates (BS, and the bootstrap-corrected (Corr.) result. Times are in units of thousands of years, effective population sizes are in thousands, and pulse amounts are in percentages.

| Run | Pop A | Pop B | $T_{ADM}$ | $T_{DIV}$ | $N_A^0$ | $N_B^0$ | $N_A^1$ | $N_B^1$ | $p$ | $N_{ANC}^0$ | $N_{ANC}^1$ |
|---|---|---|---|---|---|---|---|---|---|---|---|
| **Corr.** | **Aus.** | **Fre.** | **16.86** | **106.27** | **>1000** | **563.95** | **2.09** | **3.06** | **24.99** | **1.39** | **21.89** |
| Raw | Aus. | Fre. | 9.70 | 85.84 | 233.20 | 102.43 | 1.72 | 2.68 | 17.91 | 2.46 | 25.47 |
| BS 1 | Aus. | Fre. | 5.62 | 69.49 | 22.78 | 17.36 | 1.41 | 2.35 | 12.55 | 4.37 | 29.69 |
| BS 2 | Aus. | Fre. | 5.57 | 69.94 | 13.76 | 19.39 | 1.41 | 2.36 | 12.38 | 4.44 | 29.68 |
| BS 3 | Aus. | Fre. | 5.47 | 69.91 | 12.84 | 20.18 | 1.45 | 2.33 | 12.27 | 4.30 | 29.67 |
| BS 4 | Aus. | Fre. | 5.37 | 69.23 | 11.80 | 16.03 | 1.41 | 2.34 | 12.21 | 4.37 | 29.51 |
| BS 5 | Aus. | Fre. | 5.77 | 68.75 | 11.95 | 23.22 | 1.40 | 2.30 | 12.57 | 4.40 | 29.67 |
| BS 6 | Aus. | Fre. | 5.86 | 68.44 | 10.29 | 18.18 | 1.40 | 2.33 | 12.95 | 4.42 | 29.53 |
| BS 7 | Aus. | Fre. | 5.72 | 69.76 | 12.43 | 19.47 | 1.43 | 2.36 | 12.70 | 4.29 | 29.61 |
| BS 8 | Aus. | Fre. | 5.53 | 68.64 | 12.56 | 18.12 | 1.42 | 2.32 | 12.40 | 4.36 | 29.64 |
| BS 9 | Aus. | Fre. | 5.40 | 69.87 | 9.36 | 18.38 | 1.42 | 2.35 | 12.33 | 4.29 | 29.70 |
| BS 10 | Aus. | Fre. | 5.49 | 69.33 | 12.60 | 16.67 | 1.43 | 2.33 | 12.70 | 4.36 | 29.65 |
| **Corr.** | **Aus.** | **Han** | **11.28** | **91.22** | **>1000** | **>1000** | **2.16** | **2.60** | **24.53** | **1.42** | **21.49** |
| Raw | Aus. | Han | 7.68 | 78.23 | 243.41 | >1000 | 1.77 | 2.27 | 18.85 | 2.30 | 25.62 |
| BS 1 | Aus. | Han | 5.23 | 67.08 | 17.24 | 28.82 | 1.47 | 1.97 | 14.73 | 3.76 | 30.51 |
| BS 2 | Aus. | Han | 5.23 | 67.08 | 12.77 | 25.71 | 1.44 | 1.99 | 14.06 | 3.73 | 30.48 |
| BS 3 | Aus. | Han | 5.23 | 67.09 | 23.02 | 34.50 | 1.46 | 1.98 | 14.31 | 3.71 | 30.54 |
| BS 4 | Aus. | Han | 5.23 | 67.08 | 13.52 | 20.25 | 1.43 | 1.99 | 14.46 | 3.78 | 30.62 |
| BS 5 | Aus. | Han | 5.23 | 67.08 | 13.19 | 27.12 | 1.44 | 1.96 | 14.20 | 3.81 | 30.42 |
| BS 6 | Aus. | Han | 5.23 | 67.08 | 17.43 | 17.38 | 1.44 | 1.99 | 13.69 | 3.71 | 30.60 |
| BS 7 | Aus. | Han | 5.23 | 67.08 | 16.92 | 36.18 | 1.46 | 1.96 | 14.31 | 3.76 | 30.60 |
| BS 8 | Aus. | Han | 5.23 | 67.08 | 15.19 | 25.50 | 1.43 | 1.99 | 14.24 | 3.75 | 30.59 |
| BS 9 | Aus. | Han | 5.23 | 67.08 | 23.67 | 22.32 | 1.44 | 1.98 | 14.50 | 3.76 | 30.52 |
| BS 10 | Aus. | Han | 5.23 | 67.08 | 18.77 | 22.97 | 1.43 | 1.98 | 13.89 | 3.72 | 30.51 |
| **Corr.** | **Aus.** | **Pap.** | **5.30** | **33.92** | **37.95** | **52.91** | **4.33** | **2.24** | **15.35** | **2.20** | **20.59** |
| Raw | Aus. | Pap. | 5.75 | 29.80 | 72.40 | 44.36 | 2.73 | 1.71 | 16.76 | 2.47 | 24.76 |
| BS 1 | Aus. | Pap. | 6.37 | 26.45 | 130.25 | 40.28 | 1.72 | 1.30 | 18.96 | 2.76 | 29.82 |
| BS 2 | Aus. | Pap. | 6.33 | 25.90 | 54.63 | 35.07 | 1.71 | 1.33 | 18.06 | 2.76 | 29.93 |
| BS 3 | Aus. | Pap. | 6.08 | 25.93 | 97.99 | 23.10 | 1.73 | 1.32 | 17.69 | 2.76 | 29.66 |
| BS 4 | Aus. | Pap. | 6.21 | 26.36 | 139.03 | 30.50 | 1.70 | 1.32 | 17.88 | 2.77 | 29.81 |
| BS 5 | Aus. | Pap. | 6.06 | 25.93 | 89.78 | 38.19 | 1.74 | 1.30 | 18.07 | 2.76 | 29.80 |
| BS 6 | Aus. | Pap. | 6.36 | 26.21 | 139.22 | 51.78 | 1.74 | 1.30 | 18.22 | 2.76 | 29.69 |
| BS 7 | Aus. | Pap. | 6.40 | 26.39 | 123.46 | 35.35 | 1.74 | 1.29 | 18.31 | 2.76 | 29.73 |
| BS 8 | Aus. | Pap. | 6.40 | 26.42 | 224.18 | 51.22 | 1.74 | 1.31 | 19.06 | 2.77 | 29.92 |
| BS 9 | Aus. | Pap. | 6.08 | 26.16 | 158.78 | 38.44 | 1.69 | 1.31 | 17.97 | 2.78 | 29.74 |
| BS 10 | Aus. | Pap. | 6.07 | 26.15 | 474.40 | 36.94 | 1.76 | 1.32 | 18.50 | 2.77 | 29.76 |
| **Corr.** | **Fre.** | **Han** | **8.23** | **53.62** | **100.82** | **>1000** | **4.11** | **3.05** | **14.78** | **2.71** | **20.96** |
| Raw | Fre. | Han | 6.92 | 44.41 | 76.45 | >1000 | 3.17 | 2.50 | 13.22 | 3.06 | 24.04 |
| BS 1 | Fre. | Han | 5.83 | 35.77 | 45.74 | 73.12 | 2.46 | 2.06 | 11.75 | 3.46 | 27.63 |
| BS 2 | Fre. | Han | 5.57 | 36.54 | 35.76 | 171.07 | 2.45 | 2.03 | 11.59 | 3.44 | 27.56 |

| | | | | | | | | | | | |
|---|---|---|---|---|---|---|---|---|---|---|---|
| BS 3 | Fre. | Han | 5.74 | 36.39 | 78.74 | 106.41 | 2.43 | 2.07 | 11.44 | 3.45 | 27.56 |
| BS 4 | Fre. | Han | 5.94 | 37.28 | 55.90 | 294.03 | 2.46 | 2.04 | 11.84 | 3.45 | 27.53 |
| BS 5 | Fre. | Han | 5.87 | 37.31 | 86.77 | 102.31 | 2.46 | 2.03 | 11.81 | 3.44 | 27.61 |
| BS 6 | Fre. | Han | 6.02 | 37.28 | 96.23 | 174.07 | 2.42 | 2.04 | 11.85 | 3.48 | 27.62 |
| BS 7 | Fre. | Han | 5.88 | 37.29 | 42.62 | 80.18 | 2.43 | 2.04 | 12.03 | 3.47 | 27.59 |
| BS 8 | Fre. | Han | 5.73 | 36.26 | 55.81 | 286.05 | 2.44 | 2.05 | 11.87 | 3.49 | 27.53 |
| BS 9 | Fre. | Han | 5.89 | 36.26 | 45.09 | 162.32 | 2.42 | 2.04 | 11.83 | 3.45 | 27.60 |
| BS 10 | Fre. | Han | 5.72 | 37.48 | 66.54 | 122.91 | 2.45 | 2.05 | 11.98 | 3.45 | 27.57 |
| **Corr.** | **Fre.** | **Pap.** | **20.79** | **106.04** | **439.57** | **927.98** | **3.24** | **1.79** | **23.49** | **0.74** | **22.17** |
| Raw | Fre. | Pap. | 12.69 | 93.29 | 103.86 | 102.28 | 2.76 | 1.59 | 16.54 | 1.98 | 26.18 |
| BS 1 | Fre. | Pap. | 7.87 | 83.50 | 23.27 | 10.30 | 2.35 | 1.43 | 11.47 | 5.35 | 31.00 |
| BS 2 | Fre. | Pap. | 7.51 | 81.93 | 18.94 | 10.18 | 2.35 | 1.43 | 11.25 | 5.35 | 30.91 |
| BS 3 | Fre. | Pap. | 7.72 | 82.59 | 34.28 | 16.10 | 2.36 | 1.43 | 11.37 | 5.44 | 30.90 |
| BS 4 | Fre. | Pap. | 7.77 | 81.20 | 32.06 | 12.85 | 2.33 | 1.42 | 11.24 | 5.41 | 30.83 |
| BS 5 | Fre. | Pap. | 7.77 | 82.01 | 22.12 | 9.96 | 2.36 | 1.42 | 11.38 | 5.33 | 30.99 |
| BS 6 | Fre. | Pap. | 7.83 | 82.18 | 26.32 | 11.86 | 2.37 | 1.42 | 11.33 | 5.17 | 30.95 |
| BS 7 | Fre. | Pap. | 7.70 | 82.32 | 18.23 | 9.39 | 2.34 | 1.43 | 11.11 | 5.20 | 30.84 |
| BS 8 | Fre. | Pap. | 7.64 | 81.22 | 23.34 | 10.31 | 2.33 | 1.42 | 11.20 | 5.24 | 30.88 |
| BS 9 | Fre. | Pap. | 7.86 | 81.76 | 25.50 | 10.85 | 2.33 | 1.43 | 11.60 | 5.39 | 30.85 |
| BS 10 | Fre. | Pap. | 7.78 | 82.05 | 25.89 | 12.33 | 2.35 | 1.40 | 11.46 | 5.40 | 31.04 |
| **Corr.** | **Han** | **Pap.** | **18.59** | **112.60** | **>1000** | **747.50** | **2.84** | **1.82** | **26.33** | **0.92** | **22.03** |
| Raw | Han | Pap. | 10.54 | 87.92 | >1000 | 92.26 | 2.35 | 1.59 | 17.54 | 2.05 | 26.35 |
| BS 1 | Han | Pap. | 6.19 | 69.01 | 31.79 | 12.33 | 1.93 | 1.39 | 11.44 | 4.55 | 31.38 |
| BS 2 | Han | Pap. | 6.01 | 69.60 | 28.05 | 10.94 | 1.96 | 1.37 | 11.21 | 4.58 | 31.60 |
| BS 3 | Han | Pap. | 5.85 | 69.16 | 20.66 | 10.21 | 1.94 | 1.39 | 11.40 | 4.50 | 31.48 |
| BS 4 | Han | Pap. | 6.19 | 67.98 | 34.17 | 14.18 | 1.95 | 1.37 | 11.72 | 4.56 | 31.58 |
| BS 5 | Han | Pap. | 5.85 | 68.67 | 29.81 | 9.62 | 1.95 | 1.39 | 11.11 | 4.56 | 31.42 |
| BS 6 | Han | Pap. | 5.86 | 67.35 | 27.81 | 12.71 | 1.93 | 1.41 | 10.84 | 4.57 | 31.43 |
| BS 7 | Han | Pap. | 5.65 | 69.30 | 28.02 | 10.01 | 1.93 | 1.41 | 10.49 | 4.47 | 31.47 |
| BS 8 | Han | Pap. | 6.08 | 68.24 | 53.42 | 10.90 | 1.92 | 1.38 | 11.54 | 4.67 | 31.69 |
| BS 9 | Han | Pap. | 6.01 | 68.07 | 45.32 | 11.34 | 1.93 | 1.40 | 11.10 | 4.57 | 31.42 |
| BS 10 | Han | Pap. | 6.10 | 69.15 | 33.54 | 12.41 | 1.96 | 1.37 | 11.50 | 4.56 | 31.57 |
